# XChrom: a cross-cell chromatin accessibility prediction model integrating genomic sequence and cellular context

**DOI:** 10.1101/2025.11.18.689149

**Authors:** Yuanyuan Miao, Xuan Liang, Wenwen Zhang, Dongmei Han, Yurun Li, Zhen Wang

## Abstract

Single-cell chromatin accessibility offers unique insights into transcriptional regulation beyond gene expression. However, paired datasets of these two modalities remain relatively scarce, and existing computational models cannot simultaneously predict chromatin accessibility for unseen genomic regions and cells. Here, we present XChrom, a deep learning framework for genome-wide cross-cell chromatin accessibility prediction that integrates genomic sequence and single-cell transcriptomics-derived cell identity into a unified model. Comprehensive evaluations show that XChrom consistently outperforms existing methods in cross-region and cross-cell predictions, and uniquely enables cross-both predictions. It also supports cross-sample prediction when batch effects are properly corrected. Cross-species analyses demonstrate XChrom’s robust performance and its ability to capture evolutionarily conserved regulatory rules. Applied to peripheral blood mononuclear cells from COVID-19 patients, XChrom accurately identifies transcription factor activities in novel cell subpopulations and reveals pathogenesis-associated regulatory alterations. XChrom is available as an open-source Python package, facilitating single-cell transcriptional regulation research.

## Introduction

The rapid advancement of single-cell sequencing technologies has greatly improved our ability to dissect cellular heterogeneity at single-cell resolution. For instance, single-cell RNA sequencing (scRNA-seq) has been widely used to identify novel cell subpopulations and reveal the diversity of cellular identities and functions. Nevertheless, despite its strengths, scRNA-seq alone remains insufficient to uncover the transcriptional regulatory mechanisms underlying gene expression programs ^1^. To address this gap, it is essential to consider chromatin accessibility, which is a key layer of epigenetic regulation and provides critical insights into gene regulatory dynamics. In this context, single-cell assay for transposase-accessible chromatin with sequencing (scATAC-seq) profiles genome-wide open chromatin regions at single-cell resolution, offering a complementary view of regulatory heterogeneity and cell state transitions ^2^.

Although experimental techniques for simultaneously profiling RNA and ATAC modalities in a single cell have become feasible ^3^ (e.g., 10x Multiome), compared with scRNA-seq, such multimodal datasets remain relatively limited due to the constraint of experimental costs. Several computational approaches have been developed to generate scATAC-seq profiles ^4^, which can be broadly categorized into two approaches. The first class of methods leverages established mappings between RNA and ATAC modalities. For instance, Seurat ^5^ projects unimodal scRNA-seq data into a multimodal embedding space derived from multi-omics references, identifies *k*-nearest neighbors (KNNs), and predicts chromatin accessibility by transferring ATAC profiles from the neighbors. Methods such as LS_Lab ^6^ and MultiVI ^7^ instead learn a shared latent space from multi-omics reference data and train a joint representation model to reconstruct each modality, thereby enabling decoding from RNA to ATAC signals. Although these approaches have shown success in cell embeddings and data imputation, a key limitation is that they fail to exploit the underlying DNA sequence information of open chromatin regions. As a result, they are insufficient to capture sequence-driven transcriptional regulatory relationships. It is well established that lineage-specific transcriptional regulation is primarily determined by regulatory DNA sequences ^8^. Furthermore, these models are constrained to predicting chromatin accessibility only within the predefined genomic regions included in the training data, which limits their applicability for predicting entirely novel sequences.

The second class of methods focuses on sequence-based modeling. scBasset ^9^ represents a notable example, employing a multi-task convolutional neural network (CNN) to predict binary chromatin accessibility across multiple cells. scBasset adopts the architectures of sequence-to-function models used for bulk sequencing data, which have been shown to be effective in elucidating the effects of sequence variations and uncovering regulatory rules ^10^. Although scBasset captures *cis*-regulatory features encoded in DNA sequences and performs well in cell clustering and data denoising, it cannot generalize to unseen cells, limiting its practical applicability to new samples or cell states.

To overcome these limitations, we developed XChrom, a multimodal deep learning framework for predicting chromatin accessibility from sequences across single cells. It builds upon the CNN architecture of scBasset ^9^ to model DNA sequences, effectively capturing the regulatory information encoded in DNA. Importantly, it incorporates cell identity information as a key innovation, represented as low-dimensional embeddings derived from scRNA-seq data, into the model input. Our motivation stems from the fact that RNA-derived features, such as the expression levels of key genes ^11,12^ or binding status of core transcription factors (TFs) ^13,14^, can serve as proxies for tissue or cell-line identity, thereby enabling cross-context generalization. We extend this concept to the single-cell level, leveraging the abundance of scRNA-seq data across diverse biological and disease contexts. This framework enables predicting chromatin accessibility for any genomic sequence in cells that are similar but not identical to those in the training set. Comprehensive benchmarking demonstrates that XChrom achieves excellent performance in both within-sample and cross-sample prediction scenarios. More importantly, through interpretability analyses, XChrom can reveal *cis*- and *trans*-regulatory mechanisms across varying cellular contexts. In cross-species prediction, it exhibits exceptional generalization by capturing evolutionarily conserved regulatory rules. For peripheral blood mononuclear cells (PBMCs) from COVID-19 patients, XChrom accurately captures TF activities of a cell subpopulation previously absent from the training set, and reveals dynamic TF activities associated with disease progression. Together, these results establish XChrom as a robust and generalizable framework, enabling exploration of epigenetic regulation in development and disease directly from transcriptomic data. XChrom is implemented as an open-source Python package, with detailed tutorials freely available at https://xchrom.readthedocs.io/en/latest/.

## Results

### Overview of XChrom

XChrom is a deep learning model that integrates sequence information and cell identity to predict chromatin accessibility probabilities for each genomic region across all cells. It is trained on paired scATAC-seq and scRNA-seq data from the same cells. The core design is based on the premise that the accessibility potential of genomic regions is determined by both sequence content and cellular context. Specifically, XChrom is structured as a hybrid neural network (Fig. 1a). The first component being a sequence encoding module using CNN. This module takes a 1344-bp sequence centered around each peak from scATAC-seq data as input. After one-hot encoding, the sequence is processed through seven convolutional blocks, each containing a one-dimensional convolutional layer, a batch normalization layer, a max pooling layer, and a Gaussian error linear unit (GELU) activation function. Finally, an additional convolutional layer and a fully connected layer convert each input sequence into a 32-dimensional peak embedding. The second component is a cell identity encoding module. As input, XChrom receives cell embeddings computed from scRNA-seq data of the paired dataset (e.g., embeddings from principal component analysis (PCA)). Following *Z*-score normalization and layer normalization, the initial cell embedding matrix is transformed through two fully connected layers, producing a final 32-dimensional cell embedding matrix. The peak embedding and cell embedding matrix are integrated via matrix multiplication, generating predicted chromatin accessibility probabilities for a given genomic sequence across all cells (Fig. 1a). The model is trained using the standard binary cross-entropy loss function and optimized using the Adam optimizer (Methods).

**Fig. 1.**
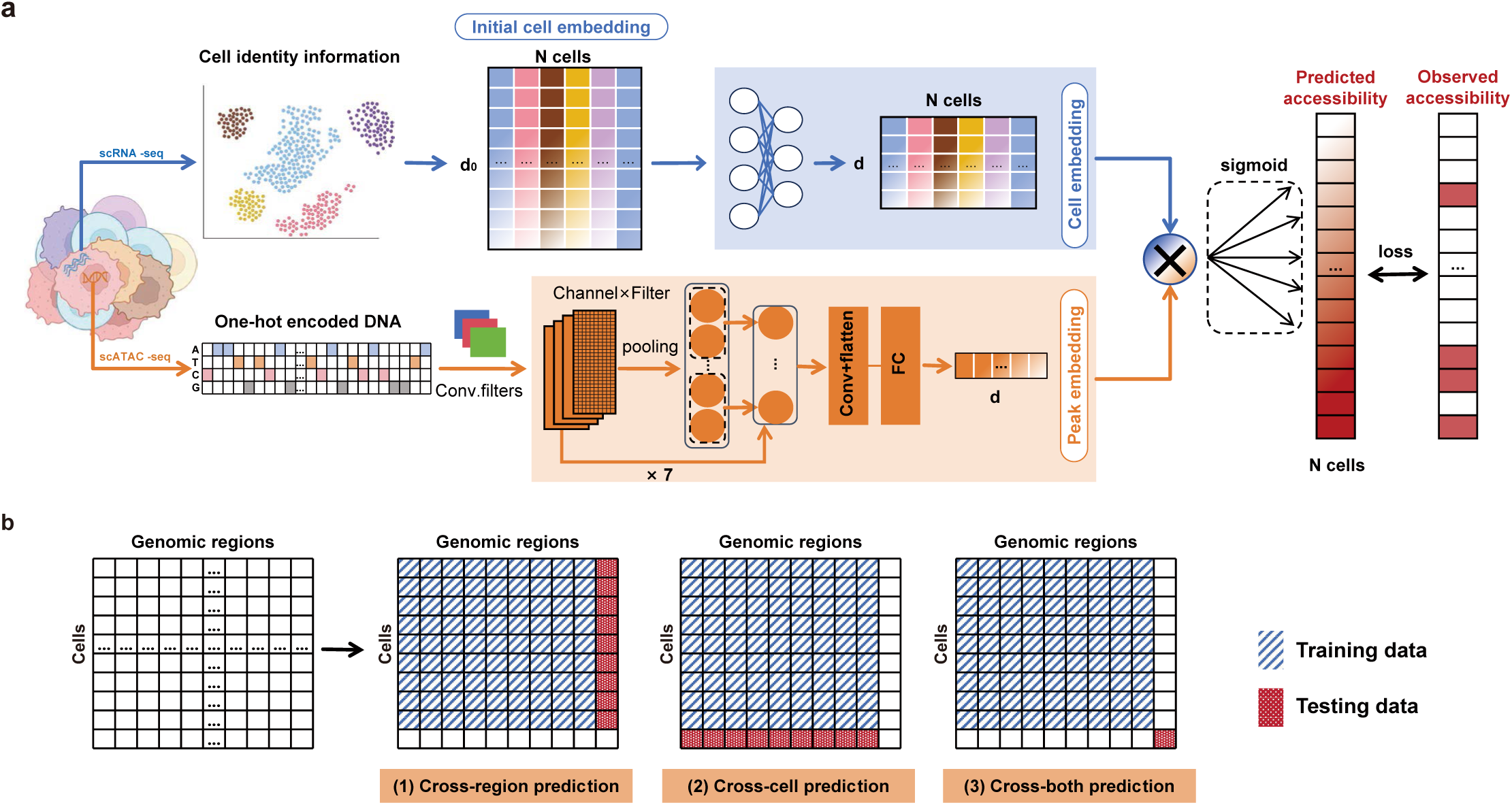
XChrom architecture and test tasks. a, XChrom is a hybrid neural network with two distinct inputs. The first input is the one-hot encoding of peak sequences (1344bp) from scATAC-seq data, processed through multi-layer convolutions to ultimately generate a peak embedding for each sequence. The second input is an initial cell embedding matrix extracted from paired scRNA-seq data, which is then transformed through dense layers to yield the final cell embedding matrix. These two sets of embeddings are integrated using matrix multiplication to predict the chromatin accessibility probabilities of a given sequence across all cells. “Conv” refers to convolutional layers, and “FC” refers to fully connected layers. *d* represents the dimensionality of the final peak and cell embeddings (default 32), while *d*_0_ represents the dimensionality of the initial cell embedding. b, The three test tasks for XChrom: the cross-region prediction, testing new regions not included in the training set; the cross-cell prediction, testing new cells not included in the training set; and the cross-both prediction, testing both unseen cells and unseen regions.

To comprehensively assess the predictive performance of XChrom, we divided the test set into three parts (Fig. 1b): (i) the “cross-region” test set, which evaluates the model’s ability to predict new genomic regions not seen during training; (ii) the “cross-cell” test set, which assesses the model’s generalization to new cells not included in the training set; and (iii) the “cross-both” test set, which tests prediction of unseen genomic regions and unseen cells, represents the most challenging task. For each test, we used two binary classification metrics-the area under the receiver operating characteristic curve (auROC) and the area under the precision-recall curve (auPRC)-as primary evaluation metrics, which were summarized at the overall, per-cell, and per-peak levels (Methods).

### XChrom accurately predicts chromatin accessibility across three test tasks

We collected five publicly available paired scRNA-seq and scATAC-seq datasets for benchmarking against existing models, covering both human and mouse species (Fig. 2a). Each dataset underwent consistent preprocessing, training/validation/test set division, and model training (Methods). The model converged after 1000 epochs of training across all datasets, as evidenced by validation set metrics (Fig. S1). In the cross-region prediction test, we compared XChrom with scBasset ^9^, a state-of-the-art chromatin accessibility prediction model that relies solely on sequence information. Although performance varied across datasets, XChrom’s performance trend was highly consistent with that of scBasset across all evaluation metrics, including overall, per-cell, and per-peak auROC/auPRC (Fig. 2b and Fig. S2, Supplementary Tables 1). This indicates that XChrom, as a multimodal model, retains the strengths of sequence models in cross-region prediction.

**Fig. 2.**
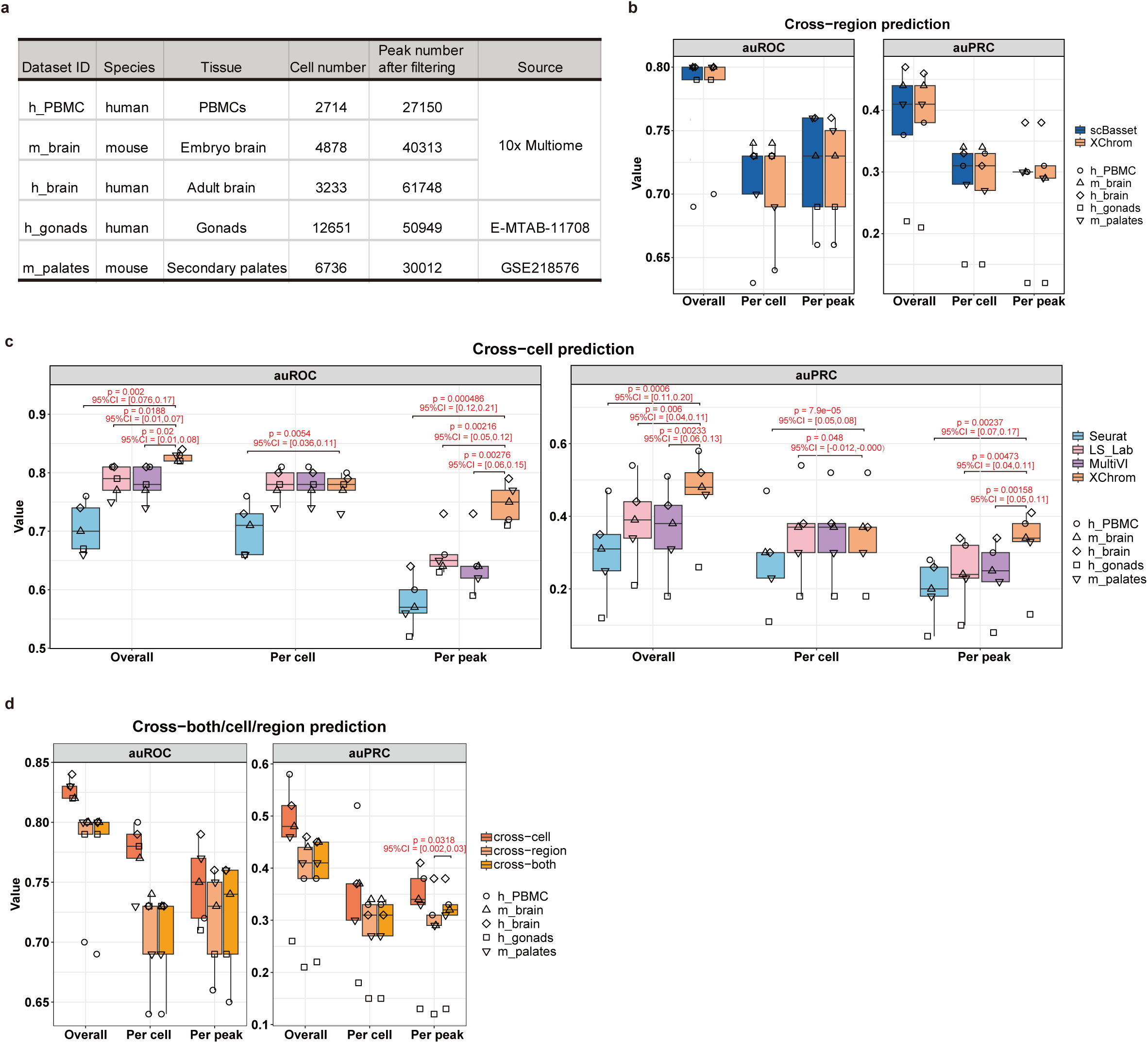
XChrom’s classification performance and benchmarking against other methods across three test tasks. a, Basic information of the publicly available paired scRNA-seq and scATAC-seq datasets used for benchmarking. b, Comparison of prediction performance between XChrom and scBasset in the cross-region prediction test. No significant differences were observed between the two models across all evaluation metrics (two-sided paired *t*-test). c, Comparison of XChrom with Seurat, LS_Lab, and MultiVI in the cross-cell prediction test. *P*-values and 95% CIs are indicated in the figure if *P*-value<0.05 (two-sided paired *t*-test). d, Evaluation of XChrom’s performance across three test tasks: cross-both, cross-region, and cross-cell. *P*-values and 95% CIs are indicated in the figure if *P*-value<0.05 (two-sided paired *t*-test).

In the cross-cell prediction test, we compared XChrom with three single-cell cross-modal prediction models: Seurat ^5^, LS_Lab ^6^, and MultiVI ^7^ (Methods). These models were top-performing in a previous benchmarking study ^4^. XChrom significantly outperformed all other models in overall and per-peak auROC/auPRC (two-sided paired *t*-test, Fig. 2c, Supplementary Tables 1). For per-cell auROC/auPRC, XChrom performed comparably to LS_Lab and MultiVI, but significantly outperformed Seurat. The advantage of XChrom in per-peak metrics indicates that it is more capable of accurately capturing cell-specific information when predicting chromatin accessibility in unseen cells.

Finally, we evaluated XChrom’s performance in the cross-both prediction test, a task infeasible for all the other models. Fig. 2d shows that, although this setting is more challenging, XChrom still achieves satisfactory prediction performance. For most metrics, its performance was not significantly different from that in the cross-region and cross-cell tasks (two-sided paired *t*-test, Supplementary Tables 1). These results indicate that XChrom retains robust generalization ability when faced with unseen genomic regions and cells, demonstrating that it learns the general principle by which DNA sequences determine chromatin accessibility in a cell-specific manner.

### Chromatin accessibility data predicted by XChrom have high cell state fidelity

Due to the inherent sparsity of scATAC-seq, binarized data often contain a large number of false-negative signals. Previous studies proposed that imputed scATAC-seq data from a machine learning model can serve to remove such noise ^9,15,16^. Therefore, we further examined XChrom’s denoising capability on raw scATAC-seq data by assessing the preservation of cell identity, which we defined as cell state fidelity. It was quantified using two metrics introduced by scBasset ^9^: the neighbor score (NS), which quantifies cross-modality neighborhood concordance by comparing scATAC-seq and scRNA-seq data, and the label score (LS), which measures the consistency of cell-type labels within local neighborhoods (Methods). Both metrics depend on a parameter *k*, the number of nearest neighbors in the KNN graph. For datasets lacking prior cell annotations, we used cluster indices identified via Leiden clustering of scRNA-seq data as proxy cell type labels for LS calculation. As expected, the NS and LS derived from denoised results showed high consistency with those from final cell embeddings, providing another perspective on monitoring convergence during model training (Fig. S3, Methods).

First, we applied the trained model for each dataset to denoise cells in the whole dataset and compared the results among XChrom, scBasset, and raw scATAC-seq data. Uniform Manifold Approximation and Projection (UMAP) visualizations revealed that the denoised data generally showed clearer separation among cell types in h_PBMC (Fig. 3a) and the other four datasets (Fig. S4). In particular, across multiple neighborhood resolution value *k*, XChrom’s denoised data consistently achieved higher NS and LS than both scBasset and raw scATAC-seq data (two-sided paired *t*-test, Fig. 3b, Supplementary Tables 2), probably because XChrom incorporated cell identity information from scRNA-seq during training.

**Fig. 3.**
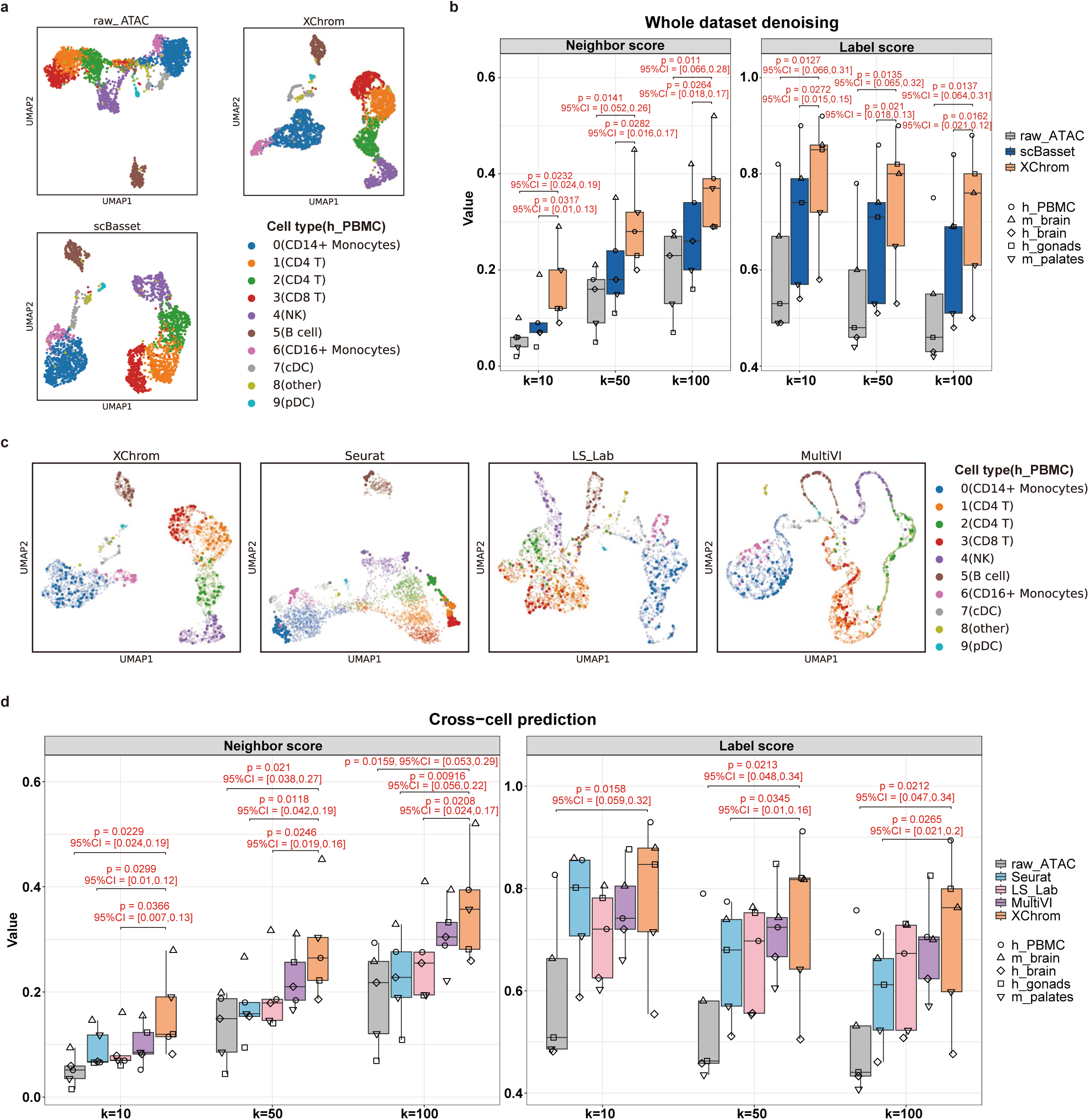
XChrom’s prediction performance in cell state fidelity and benchmarking against other methods. a, UMAP visualizations of the raw scATAC-seq data (raw_ATAC), XChrom- and scBasset-denoised results for the h_PBMC dataset, with cell type annotations for each cluster. b, NS and LS metrics computed for raw_ATAC, XChrom-, and scBasset-denoised results. Each metric was calculated with *k*=10, 50, and 100 neighbors. *P*-values and 95% CIs are indicated in the figure if *P*-value<0.05 (two-sided paired *t*-test). c, UMAP visualizations of test cell predictions by Seurat, LS_Lab, MultiVI, and XChrom for the h_PBMC dataset. Larger points represent test cells, while background points represent training cells. d, Evaluation of NS and LS for test cell predictions, comparing XChrom with Seurat, LS_Lab, MultiVI, and raw_ATAC. *P*-values and 95% CIs are shown if *P*-value<0.05 (two-sided paired *t*-test).

Subsequently, we evaluated XChrom with cross-modal prediction models, including Seurat ^5^, LS_Lab ^6^, and MultiVI ^7^, with a specific focus on the cross-cell test set for assessing cell state fidelity of unseen cells (Methods). XChrom consistently imputed correct cell labels for these test cells, as shown in UMAP visualizations (Fig. 3c, Fig. S5). When considering only NS and LS for the test cells, we found that XChrom and MultiVI performed the best in most cases (two-sided paired *t*-test, Fig. 3d, Supplementary Tables 2). However, in some instances, MultiVI’s UMAP visualizations were less than ideal, showing a linear pattern and unclear boundaries between clusters (Fig. 3c, Fig. S5).

In conclusion, these results indicate that XChrom can not only effectively denoise existing scATAC-seq data but also directly generate denoised results for cells lacking scATAC-seq data when provided with cell identity information.

### XChrom can predict new samples from the same tissue or organ as the training set

The preceding study validated the model’s performance within a single sample, where the scRNA-seq data in both the training and test sets are inherently homogeneous, without batch effects. However, in most practical scenarios, the objective is to predict completely new and independent samples from a similar tissue or organ as the training set. To address this, we further evaluated XChrom’s performance in a cross-sample prediction scenario. We collected a dataset of paired scRNA-seq and scATAC-seq from bone marrow mononuclear cells (BMMCs) ^17^, which includes multiple batches arising from donor sources and sequencing platforms. A total of five samples were randomly selected as the training set, and five as the test set, employing a one-to-one cross-sample training and testing approach. We corrected the batch effects in the scRNA-seq data from both the training and test sets using Harmony ^18^ (Fig. S6) and used the resulting cell embedding matrix as model inputs.

First, we compared the performance of cross-sample prediction to within-sample prediction by XChrom. As shown in Fig. 4a, cross-sample prediction resulted in a significant decrease in overall, per-cell, and per-peak auROC (two-sided paired *t*-test, Supplementary Tables 3). However, the 95% CIs for the differences indicated that the magnitude of the reductions was limited. In the cell state fidelity assessment on the test set, the NS and LS values for cross-sample prediction were lower than those observed in the within-sample results, but XChrom still outperformed the raw scATAC data—particularly in terms of NS, where this superiority was statistically significant (two-sided paired *t*-test, Fig. 4b, Supplementary Tables 3). These findings suggest that batch effects between samples do influence XChrom’s predictive performance, but XChrom remains applicable for predicting new samples if these batch effects are properly corrected.

**Fig. 4.**
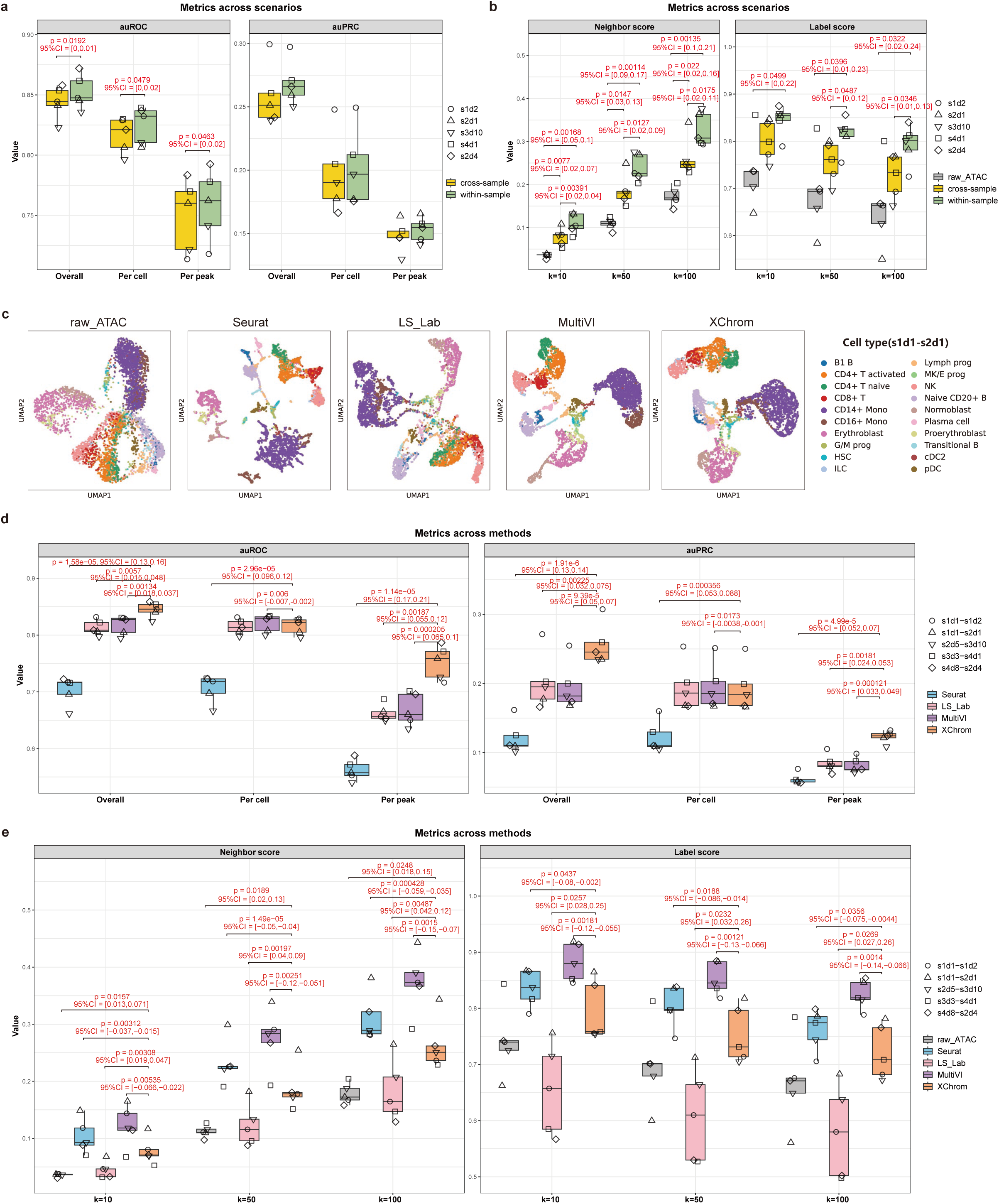
XChrom’s prediction performance and benchmarking against other methods in cross-sample scenario. a-b, Comparison of within-sample and cross-sample predictions by XChrom for classification performance (a) and cell state fidelity (b). Only sample names of the test sets are labeled. *P*-values and 95% CIs are shown in the figure if *P*-value<0.05 (two-sided paired *t*-test). c, Sample s1d1 from the BMMC dataset serving as the training set, with s2d1 as the test set. UMAP visualizations of raw scATAC-seq data (raw_ATAC) of s2d1 and predicted data of s2d1 by Seurat, LS_Lab, MultiVI, and XChrom. d-e, Comparison of raw_ATAC and cross-sample predictions by Seurat, LS_Lab, MultiVI, and XChrom for classification performance (d) and cell state fidelity (e). In the legend, sample names of ‘training set-test set’ are labelled. *P*-values and 95% CIs are indicated in the figure if *P*-value<0.05 (two-sided paired *t*-test).

We also compared XChrom with Seurat ^5^, LS_Lab ^6^, and MultiVI ^7^ in the cross-sample scenario (Fig. 4c, Fig. S7). The results demonstrated that XChrom significantly outperformed the other models across most classification metrics (two-sided paired *t*-test, Fig. 4d, Supplementary Tables 3). In terms of cell state fidelity, XChrom’s performance was at a median level compared to other models, but outperformed raw scATAC-seq data (Fig. 4e). We hypothesize that, although batch effects were corrected during preprocessing using Harmony, residual batch differences between the training and test data may slightly impair XChrom’s ability to preserve cell identity. Nonetheless, the results indicate that XChrom is adequate for predicting new samples from tissues or organs similar to those in the training set.

### XChrom enables cross-species prediction by capturing evolutionarily conserved regulatory rules

Recent studies have demonstrated that sequence models trained on mouse scATAC-seq data can accurately predict chromatin accessibility maps for other vertebrate genomes. These models have also revealed conserved regulatory logic and lineage-specific gene regulation across vertebrates ^19^. Such findings support Arendt’s hypothesis that “cell identity across species is conserved and defined by a core gene regulatory network (GRN)” ^20^. However, previous cross-species models were limited to cells from the training set. Building on XChrom’s unique cross-both prediction capacity, we examined its potential for cross-species prediction with varying genomic sequences and cellular contexts.

We collected and preprocessed paired scRNA-seq and scATAC-seq data from the primary motor cortex of four species: human, mouse, macaque, and marmoset, with each species containing at least three samples ^21^. All scRNA-seq data underwent uniform batch correction prior to training and prediction (Methods, Fig. S8). Models were trained using a single sample from human (human_model: m1d1n) and mouse (mouse_model: mop3c2), respectively. We then assessed the cross-species prediction performance of these two models on all remaining samples.

The results indicate that the predicted data from either the human_model (Fig. 5a) or mouse_model (Fig. S9a) could distinguish different cell types across each species. When applied to human test sets, the human_model and mouse_model performed comparably on most metrics, including per-peak auROC/auRPC, per-cell auROC (Fig. 5b, Fig. S9b) and NS/LS (Fig. 5c, Fig. S9c, two-sided paired *t*-test, Supplementary Tables 4). However, when applied to mouse test sets, the mouse_model significantly outperformed both the human_model and the corresponding raw scATAC-seq data across all metrics (two-sided paired t-test, Fig. 5b, c, Fig. S9b, c, Supplementary Tables 4). This could result from the fact that, as a model organism, mouse samples were more homologous with minimal potential batch effects, rather than inherent species difference. When applied to macaque and marmoset test sets, human_model and mouse_model were broadly comparable. On several metrics-including overall, per-cell, per-peak auROC/auPRC-the mouse_model even outperformed the human_model (Fig. 5b, Fig. S9b, Supplementary Tables 4). Taken together, these results suggest that XChrom effectively captured evolutionarily conserved regulatory features and principles, demonstrating robustness even across distantly related species such as primates and rodents.

**Fig. 5.**
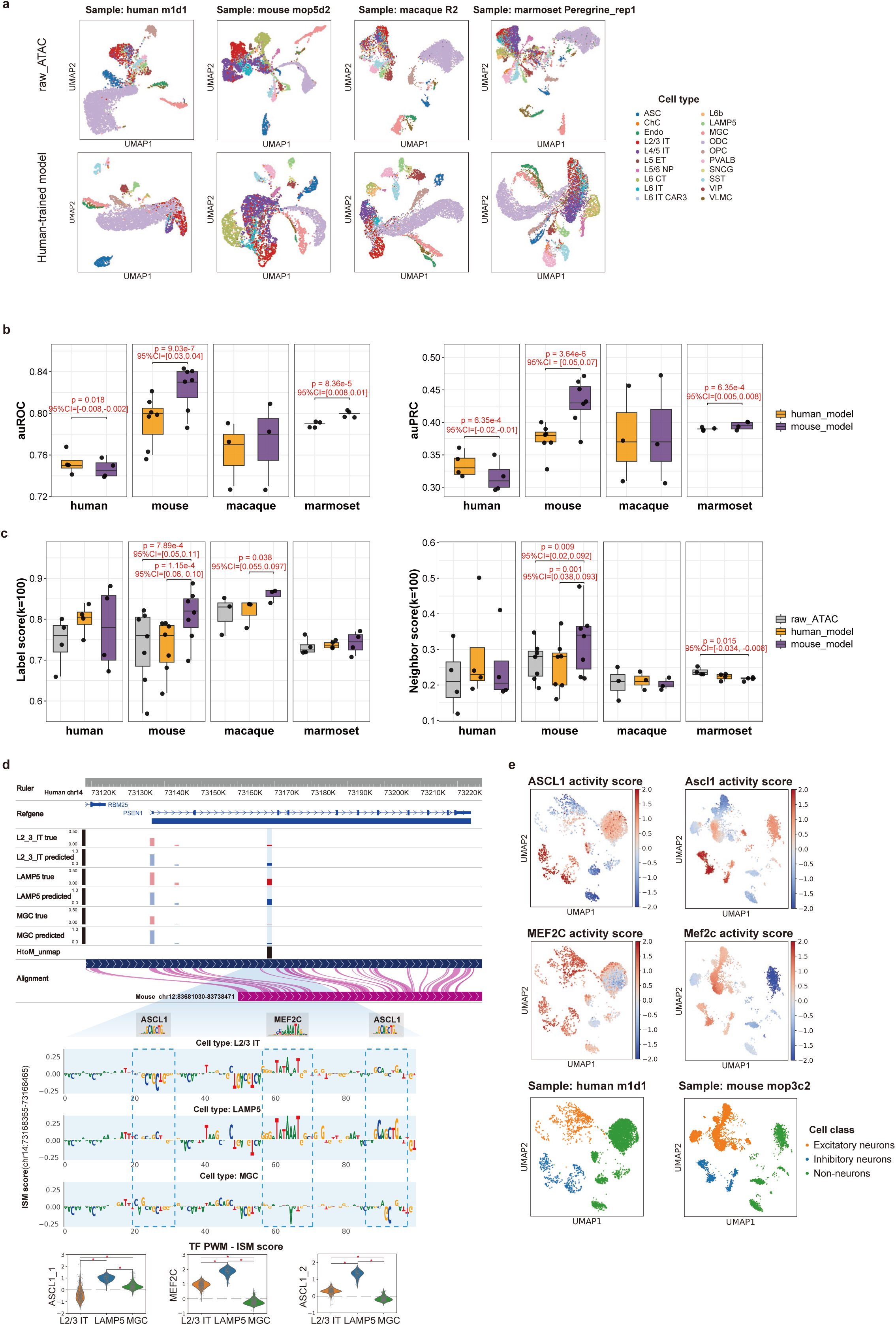
XChrom’s prediction performance in cross-species scenario and its interpretability analysis. a, UMAP visualizations of predictions generated by the human_model when applied to the test sets from four different species. Sample names are labelled in the plot. b-c, Comparison of model performance between human_model and mouse_model using auROC/auPRC values (b), and NS/LS values (*k*=100) (c), computed from predictions on all samples. *P*-values and 95% CIs are shown in the figure if *P*-value<0.05 (two-sided paired *t*-test). d, Visualization of mouse_model predictions for a *PSEN1* regulatory sequence in the human sample (m1d1), with reference data from the WashU Comparative Epigenome Browser (https://epigenome.wustl.edu/BrainComparativeEpigenome/) ^21^. The blue-shaded sequence represents the human region (chr14: 73167855-73169199) that lacks homology to the mouse genome. Subsequences matching MEF2C and ASCL1 motifs are highlighted with blue dashed boxes in the cell-type-averaged ISM score plot. Below are per-cell PWM-ISM scores for MEF2C and ASCL1 in L2/3 IT (*n*=335), LAMP5 (*n*=78), and MGC (*n*=211) cell types. The PWM-ISM score is the dot product of the PWM and ISM scores at motif match sites (MEF2C at chr14:73168422-73168436, ASCL1 at chr14:73168385-73168397 and chr14:73168451-73168463). A two-sided Wilcoxon rank-sum test was performed for statistical significance; * *P*-value<0.05. PWM: position weight matrix. e, UMAP visualizations of MEF2C and ASCL1 motif activity predicted by the mouse_model across different cell types in the human sample (left) and the mouse sample (right).

A key advantage of sequence models is their interpretability ^22^, which can be utilized to further investigate the conservation of regulatory mechanisms. We calculated the TF motif activity at the single-cell level, defined by the difference predicted by XChrom when the motif sequence was inserted into the background sequence ^23^ (Methods). We analyzed the cross-species conservation of core TF activities across cell types (Methods). The average correlation of TF activity across cell types was high between the following pairs: 1) human_model predictions for both human and mouse samples (0.61); 2) mouse_model predictions for both human and mouse samples (0.67); 3) human_model predictions for human sample and mouse_model predictions for mouse sample (0.47, Fig. S10). These results demonstrated that the conservation of cell type-specific TF activity patterns was captured by XChrom.

To verify whether the model relies solely on sequence similarity for cross-species prediction, we analyzed the neurodegenerative disease-related gene *PSEN1* ^24^. As shown in Fig. 5d (representative cell types) and Fig. S11a (other cell types), in the human sample (m1d1), a sequence that lacks homology to the mouse genome is accessible in the raw scATAC-seq data. Despite not encountering this sequence during training, the mouse_model accurately predicted its accessibility. Further in silico saturation mutagenesis (ISM) was used to evaluate the importance of each site in this sequence ^25^ across cells in the human sample. Combined with FIMO ^26^ motif scanning results (Methods), we found that binding sites of MEF2C and ASCL1 exhibited higher regulatory importance in inhibitory neurons (e.g., LAMP5) compared with excitatory neurons (e.g., L2/3 IT) and non-neurons (e.g., MGC), consistent with the higher real and predicted accessibility of this sequence in inhibitory neurons (Fig. 5d, Fig. S11a, b). These two TFs were also activated in inhibitory neurons in both human and mouse samples, as identified by XChrom’s TF activity analysis (Fig. 5e, Fig. S11c, d). Previous research has shown that MEF2C plays a key role in early brain development, with mutations linked to various neuropsychiatric disorders ^27,28^, while ASCL1 is a major regulator of neuronal fate determination during central nervous system development, particularly involved in generating inhibitory GABAergic interneurons ^29,30^.

We also analyzed another gene, *DNAJC5* (also known as *CSP*α), which plays a crucial role in synaptic vesicle recycling and neurotransmitter release ^31^. In the gene body, a human-specific sequence was identified, with higher raw and predicted accessibility in excitatory neurons. Binding sites of both TEAD2 and GLI2 showed higher prediction importance in this sequence within these cell types, with their activation observed in both human and mouse samples (Fig. S12). Previous studies have shown that TEAD2 participates in the mouse embryonic neural tube closure process ^32^, while GLI2 is closely related to neurogenesis, neuronal migration, and differentiation ^33^.

In summary, despite genomic differences between species, XChrom does not rely solely on sequence similarity for cross-species predictions. Instead, it captures conserved transcriptional regulatory mechanisms, enabling robust transfer and generalization capabilities across species.

### XChrom predicts TF activities of novel cell subpopulations and regulatory changes across different COVID-19 conditions

To explore the potential application of XChrom for new samples with different physiological and pathological conditions, we applied it to a single-cell sequencing dataset of PBMCs from COVID-19 patients, including severe, mild, and convalescent cases ^34^. Although the dataset contains both scRNA-seq and scATAC-seq data, they are not paired. Therefore, we used a public paired scRNA-seq and ATAC-seq dataset from healthy donors (PBMC10k, 10x Multiome) to train an XChrom model, and then predicted the epigenomic features of COVID-19 patients based on their scRNA-seq data. The PBMC10k and COVID-19 scRNA-seq data were harmonized prior to model training and prediction (Fig. S13a). The unpaired COVID-19 scATAC-seq data served as an independent resource for validating the model’s findings.

First, we inferred the TF activity across different cell types (Methods). The results showed that XChrom could accurately identify well-established cell type-specific TF activities, such as CEBPB (monocytes), TCF7 (T cells), PAX5 (B cells), and RUNX3 (NK cells) (Fig. S13b). As the pathogenesis of COVID-19 is highly associated with monocytes ^34^, we focused on this cell type in the following analysis (Fig. 6a). In the processed COVID-19 scRNA-seq data, a subcluster of monocytes (R1) was annotated as non-classical monocytes (ncMono). Differential analysis of TF activity in R1 compared to other subclusters revealed significantly higher activity of several TFs (e.g., IRF family members, SPI1, STAT2, two-sided Wilcoxon rank-sum test, Supplementary Tables 5), which were also significantly enriched in the corresponding subcluster (C1) in the COVID-19 scATAC-seq data (Fig. 6b, Supplementary Tables 5, Methods). These TFs are known to play key roles in myeloid development, antiviral response, and inflammation ^35–41^.

**Fig. 6.**
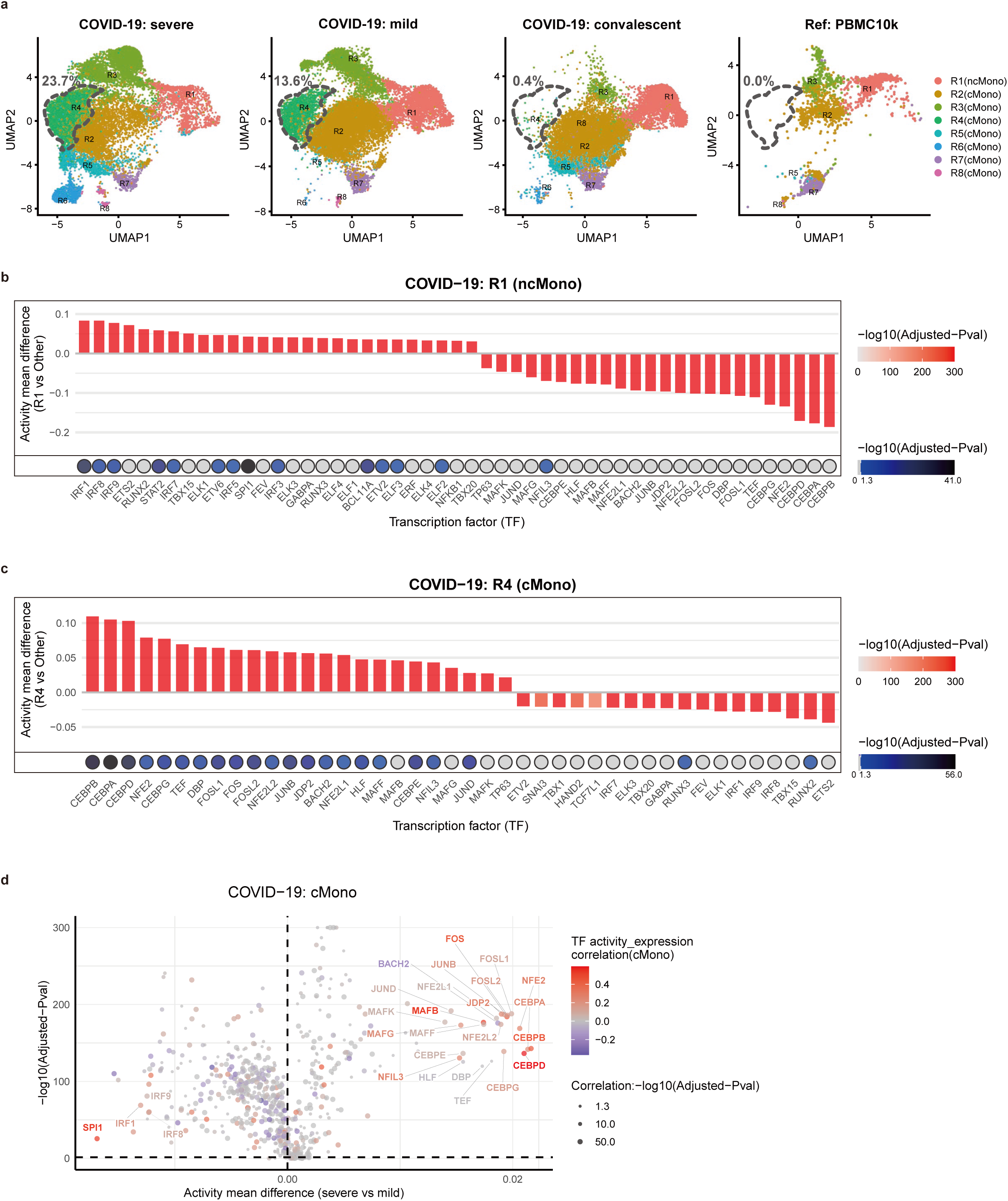
XChrom infers single-cell TF activity and enables differential analysis across different COVID-19 conditions. a, UMAP visualizations of monocytes from the COVID-19 scRNA-seq dataset (including severe, mild, and convalescent patients) and the PBMC10k scRNA-seq dataset. Subclusters of monocytes are labelled within each dataset. The grey dashed line indicates R4 subcluster and its proportion. b-c, Differential TF activity (bar plots) and motif enrichment (heatmaps) analysis in COVID-19 monocyte subclusters. Colors reflect adjusted *P*-values (two-sided Wilcoxon rank-sum test for TF activity; motif enrichment considered significant at -log10 (adjusted *P*-value) > 1.3). R1 vs. other subclusters: bar plot shows the top 51 TFs with |mean difference| > 0.03 (positive values indicate higher activity in R1); heatmap shows motifs enriched in marker peaks of the corresponding subpopulation (C1) in the COVID-19 scATAC-seq data (b). R4 vs. other subclusters: bar plot shows the top 42 TFs with |mean difference| > 0.02 (positive values indicate higher activity in R4); heatmap shows motifs enriched in marker peaks of the corresponding subpopulation (C4) in the COVID-19 scATAC-seq data (c). d, Differential analysis of TF activity in cMono between mild and severe COVID-19 patients. The x-axis represents the mean TF activity difference (severe-mild), and the y-axis represents the adjusted *P*-value of the difference (two-sided Wilcoxon rank-sum test). The color of the dots indicates the Pearson correlation between TF activity and its expression, while the size of the dots indicates the adjusted *P*-value of this correlation. The volcano plot uses a pseudo-log transformation for axis scaling.

Next, we tested XChrom’s generalization capability by predicting the epigenomic states of a novel classical monocyte (cMono) subpopulation (R4) that was absent from the training set (Fig. 6a). In fact, R4 was annotated as HLA-DR^lo^S100A^hi^ monocytes in the original study, which were mainly derived from hospitalized COVID-19 patients (both mild and severe) but absent from convalescent cases^34^. TF activity prediction by XChrom and subsequent differential analysis on R4 revealed that C/EBP, FOS, and JUN family members were significantly more active in R4 than in other subclusters (two-sided Wilcoxon rank-sum test, Supplementary Tables 5). These TFs were also significantly enriched in the corresponding subpopulation (C4) in the COVID-19 scATAC-seq data (Fig. 6c, Supplementary Tables 5), and this was consistent with the study showing that the dysfunctional HLA-DR^lo^S100A^hi^ monocytes were characterized by CEBPD, CEBPE, FOSL2, FOSB, JUN, and NFIL3 in severe COVID-19 patients ^42^. These results indicate that XChrom can not only effectively recover the regulatory features of cell types contained in the training set, but also learn to extrapolate the regulatory logic to novel cell subpopulations.

Finally, we investigated regulatory changes during the progression of COVID-19 by comparing the predicted TF activities in cMono between mild and severe COVID-19 patients (Methods). We found that TFs such as C/EBP, FOS, JUN, and MAFB exhibited stronger activity in severe cases (two-sided Wilcoxon rank-sum test, Supplementary Tables 5), with their activity often positively correlating with gene expression (Fig. 6d). These TFs are closely linked to the exacerbated inflammatory response and pro-fibrotic processes observed in severe COVID-19: activation of C/EBP TFs has been reported to promote profibrotic macrophage differentiation and trigger lung fibrosis in COVID-19 ^43^. FOSL1 has been shown to regulate the expression of pro-inflammatory/anti-inflammatory cytokines in macrophages and modulate pro-fibrotic responses^44^. MAFB is a key driver in the transition from a macrophage-to-inflammatory-fibrotic pathological state following SARS-CoV-2 infection, and its silencing significantly attenuates the associated pathogenic phenotype ^45^. The balance between MAFB and MAF in lung macrophages is a critical determinant of COVID-19 severity, with increased MAFB/MAF ratios weakening the type I interferon response, exacerbating inflammation, and enhancing fibrosis ^46^. Additionally, MAFB can form heterodimers with JUN, FOS, and FOSL1/2 members of the AP-1 family ^47^. AP-1, as a key downstream effector of the MAPK signaling pathway, is closely related to macrophage inflammatory responses induced by pathogen-associated molecular patterns and is involved in the “cytokine storm” observed in viral infections ^48,49^. In contrast, the activities of IRF family members (IRF1/8/9) were significantly reduced in severe cases (two-sided Wilcoxon rank-sum test, Fig. 6d, Supplementary Tables 5), which may impair the expression of interferons with antiviral roles. In fact, deficiency in type I interferon production and activity was observed in severe COVID-19 patients ^50^, and a low-interferon cMono state was predominant in severe cases ^51^. Together, these results suggest that XChrom can make biologically plausible TF activity predictions across disease conditions.

## Discussion

In this study, we present XChrom, a deep learning-based framework capable of predicting genome-wide chromatin accessibility at single-cell resolution. The core innovation of XChrom lies in integrating DNA sequence information with cell identity derived from scRNA-seq into a unified neural network architecture. Capturing the dependence between *cis*-regulatory mechanisms and cellular context allows XChrom to overcome the limitations of existing methods, thereby enabling accurate inference of chromatin accessibility in both unseen sequences and unseen cells.

Through comprehensive evaluations, XChrom demonstrated superior predictive performance across various scenarios. Specifically, in cross-region prediction within a sample, XChrom achieved comparable classification performance to scBasset ^9^ while showing better noise reduction capability. In cross-cell prediction within a sample, XChrom outperformed Seurat ^5^, LS_Lab ^6^, and MultiVI ^7^ in classification performance, while maintaining the highest cell identity fidelity along with MultiVI. Additionally, XChrom was able to predict the accessibility probabilities of unseen genomic regions in entirely unseen cells, a task that existing models fail to accomplish. In cross-sample prediction, although XChrom was influenced by batch effects in scRNA-seq data between training and test samples, it still achieved optimal classification performance. Its performance in preserving cell identity was intermediate compared to Seurat, LS_Lab, and MultiVI, but consistently outperformed raw scATAC-seq data.

Additionally, XChrom exhibits strong interpretability, facilitating understanding of both *cis*- and *trans*-regulatory mechanisms at single-cell resolution. For local interpretation, through ISM analysis, XChrom enables assessment of the contribution of individual site within a *cis*-regulatory element to chromatin accessibility, offering a new approach for prioritizing non-coding genetic variants. For global interpretation, by inserting TF motif sequences into background sequences, XChrom can infer TF motif activity at the single-cell level, providing a new perspective to study cell type-specific *trans*-regulatory mechanisms beyond gene expression.

These features enable us to apply XChrom to entirely new scenarios. Using a paired scRNA-seq and scATAC-seq dataset from the primary motor cortex of four species, we demonstrated that XChrom trained on human and mouse data achieved excellent prediction performance in other species as well. In some cases, the performance of cross-species predictions was even comparable to that of within-species predictions. Our local interpretability analyses of chromatin accessible regions lacking homology across species revealed that XChrom does not rely on simple sequence similarity; instead, it captures evolutionarily conserved regulatory rules, such as binding sites of key TFs. These results indicate that we can train XChrom in model organisms with abundant single-cell data and then apply it to non-model organisms with limited data to study their genetics and evolution.

In the case study of PBMC data from COVID-19 patients, XChrom not only accurately predicted TF activities of a novel cell subpopulation that was absent from the training set, but also successfully captured the dynamic changes in TF activity during the transition from mild to severe states in cMono cells. These findings indicate that XChrom does not simply memorize shallow RNA-to-ATAC mappings from the training data but instead learns the underlying biological principles linking cell states to chromatin accessibility. As a result, this enables XChrom to leverage readily available scRNA-seq data to predict epigenomic features of samples with physiological or pathological conditions distinct from those in the training set, and to explore regulatory alterations underlying these conditions.

However, it is important to note that XChrom currently has some limitations. First, the model does not explicitly account for genomic variation between samples (such as single-nucleotide polymorphisms, copy number variations, etc.), an important source of differences in sequence regulatory activity. Second, when assessing the effects of genetic variations or TF motif insertions, we assume that the cell identity remains constant, i.e., the peak embeddings and cell embeddings are independent. However, there may be a coupling between these two embeddings. Third, although XChrom can identify the activity of key TFs, the current model architecture does not directly capture the relationships between these TFs and their target genes, making it challenging to construct complete GRNs at the single-cell level.

In the future, with increased computational resources, advanced foundation models or pretrained models can be leveraged to improve data representations—for example, Geneformer ^52^ and scGPT ^53^ for cell embeddings, and Nucleotide Transformer ^54^ and Sei ^55^ for peak embeddings. This may better address batch effects, improve the recognition of rare and complex cell states, and enhance the accuracy of perturbation predictions.

## Methods

### XChrom architecture

XChrom employs a hybrid neural network architecture to predict chromatin accessibility probabilities across genomic regions at single-cell resolution (Fig. 1a). The model is built on the TensorFlow (v.2.6.0) framework and consists of two primary components:

#### i Genomic sequence encoding module

This module is based on a CNN architecture, inspired by the deep learning framework of scBasset ^9^. The input consists of DNA sequences centered around each peak defined by scATAC-seq, with a length of 1344 bp. After one-hot encoding, these sequences are processed through seven convolutional blocks. Each block includes a one-dimensional convolutional layer, a batch normalization layer, a max pooling layer, and a GELU activation function. The output dimensions at each convolutional block are as follows: Initial input (1344, 4) → □ (448, 288) → □ (224, 288) → □ (112, 323) → □ (56, 363) → □ (28, 407) → □ (14, 456) → □ (7, 512). Then, the feature map passes through an additional convolutional layer with a GELU activation, resulting in a dimension of (7, 256). Finally, the data flows through a fully connected layer (input dimension 1792, output dimension 32), producing a 32-dimensional embedding for each peak region. A dropout rate of 0.2 is used in the fully connected layer to regularize the model, randomly masking neurons to improve generalization. Additionally, sequence inputs are augmented using stochastic reverse complementation (randomly flipping strand orientation with 50% probability) and random shifts (±3 bp) as implemented in scBasset. These augmentations promote strand and positional invariance during training.

#### ii Cell identity encoding module

This module extracts cell identity information from scRNA-seq data. The initial cell embedding matrix (e.g., PCA-processed) is layer-normalized to eliminate variance contribution differences among principal components. The normalized matrix is then processed through two fully connected layers (with 64 and 32 neurons, respectively), using ReLU and linear activation functions, producing a 32-dimensional cell embedding matrix. To enhance the model’s robustness, a data augmentation strategy is applied: isotropic Gaussian noise (with a standard deviation controlled by the hyperparameter noise factor, default 0.01) is added exclusively during training. During evaluation, the input remains unchanged. This operation is implemented with the tensorflow.keras.backend.in_train_phase() function.

The 32-dimensional peak embedding and cell embedding matrix are combined through matrix multiplication. The result is then passed through a Sigmoid activation function, yielding continuous chromatin accessibility probabilities in the range [0, 1] across all cells.

Notably, XChrom explicitly considers the impact of single-cell sequencing depth during prediction. After embedding integration, a learnable sequencing depth correction term is introduced for each cell. This term is implemented through a single-neuron linear fully connected layer, allowing the model to automatically learn and adjust for sequencing depth effects on predictions. The initial input of the correction term is the sequencing depth estimated from binarized scATAC-seq data and normalized across cells by *Z*-scores. When generating the sequencing depth-normalized chromatin accessibility profiles, the default input of the correction term for all cells is set to 1.

### Model evaluation metrics

Since XChrom predicts binarized chromatin accessibility (with sequencing count ≥ 1 as the true label), auROC and auPRC are the primary evaluation metrics for the model’s discriminative ability. While auROC is relatively insensitive to class imbalance, auPRC is a more informative metric for highly imbalanced datasets. These metrics were calculated at three levels:

(i) overall metrics: calculated across all test cells and peak regions;
(ii) per-peak metrics: calculated for each peak region across test cells, then averaged over all test peak regions;
(iii) per-cell metrics: calculated for each cell across all test peak regions, then averaged over all test cells.

We also adopted NS and LS to evaluate the cell identity fidelity of raw and denoised scATAC-seq data. Both metrics were calculated using KNN graphs at different neighborhood scales (*k*=10, 50, 100), which were constructed in PCA-reduced space.

(i) NS: Independent KNN graphs were constructed using paired scRNA-seq and scATAC-seq data, respectively. NS quantifies the percentage of overlapping neighbors for each cell between the two graphs.
(ii) LS: For a given KNN graph of scATAC-seq data, LS quantifies the percentage of a cell’s neighbors that share the same cell-type label within the neighborhood. Since ground-truth cell types were unavailable for some datasets, Leiden clusters derived from scRNA-seq data were used as proxy cell-type labels.

### XChrom training strategy

Model parameters were updated using the Adam optimization algorithm, with hyperparameters set to β□ = 0.95 and β□ = 0.9995. The learning rate was decayed exponentially, starting at 0.01 and multiplied by a decay factor of 0.9 after every 10,000 training steps. The loss function used was binary cross-entropy, with binary classification accuracy, auROC, and auPRC monitored simultaneously as evaluation metrics during training. Early stopping was employed, with training terminated if the improvement in auROC on the training set was less than 1×10□□ over 50 consecutive training epochs.

The default maximum number of training epochs was set to 1000. This choice was based on an analysis of the h_PBMC and m_brain datasets: During training, we saved model parameters every 10 epochs and found that cell state fidelity metrics (NS(*k*=100) and LS(*k*=100)), calculated from the final cell embeddings, were highly correlated with the same metrics calculated after denoising the entire dataset (Fig. S3a). Consequently, we added the NS(*k*=100) and LS(*k*=100) metrics, computed from the final cell embeddings, to monitor the model’s ability to preserve cell state during training. These metrics increased steadily with more epochs and plateaued around 1000 epochs (Fig. S3b). In contrast, traditional training/validation loss and training/validation auROC/auPRC metrics stabilized early (within 250 epochs) (Fig. S1).

### Paired scRNA-seq and scATAC-seq dataset preprocessing

We collected multiple paired scRNA-seq and scATAC-seq datasets for model training and testing. We performed quality control (QC) on these datasets using scanpy (v.1.9.5) ^56^. Specifically, for scRNA-seq data, we excluded genes expressed in fewer than 5% of cells; for scATAC-seq data, we excluded peaks present in fewer than 5% of cells. For the h_gonads dataset and the cross-sample BMMC datasets, the threshold was adjusted to 1%. Additionally, we filtered out peaks not mapped to chromosomes: for human data, peaks on chromosomes 1–22, X, and Y were retained; for mouse data, peaks on chromosomes 1–19, X, and Y were retained.

### Batch effect correction in scRNA-seq data

To eliminate potential technical variations among different samples or species, we performed systematic batch effect correction on the scRNA-seq data using scanpy (v.1.9.5) ^56^. For different samples of the same species, we first integrated the scRNA-seq data based on shared genes. The integrated expression matrix was then subjected to PCA (via scanpy.pp.pca() function with n_comps=32). Next, we applied the Harmony ^18^ algorithm (via scanpy.external.pp.harmony_integrate() function), using sample batch as the correction variable.

In the cross-species scenario, we first concatenated samples within each species based on shared genes. Then, we obtained a one-to-one ortholog gene list between each species and human via BioMart ^57^, and used this list to retain orthologous genes for cross-species data integration. Each sample was treated as an independent batch, and the Harmony ^18^ algorithm was applied for batch correction across species.

After correction, data for each sample was saved as an h5ad file, with the corrected cell embedding matrix stored in .obsm[’X_pca_harmony’], which was used as the initial cell embedding matrix for model training and prediction.

### Training/test set split

To systematically evaluate model performance, we adopted the following strategies for splitting the training and test sets across different scenarios. In the within-sample scenario, we divided the cells and peak regions at a ratio of 90%, constructing four data sets for model training and evaluation (Fig. 1b).

(i) Training-validation set: composed of training cells and training peaks, with 90% of peaks used for model parameter training and 10% reserved for validation.
(ii) Cross-region test set: composed of training cells and testing peaks, used to assess the model’s ability to generalize to unseen genomic regions.
(iii) Cross-cell test set: composed of testing cells and training peaks, used to evaluate the model’s prediction performance on unseen cells.
(iv) Cross-both test set: composed of testing cells and testing peaks, used to assess the model’s overall generalization ability on unseen cells and genomic regions.

In the cross-sample or cross-species scenario, we divided the cells and peak regions of one sample at a 90% ratio for the training-validation set, and the remaining 10% served as the within-sample or within-species test set. We used another independent sample as the cross-sample or cross-species test set to evaluate the model’s transferability to different samples or species.

### Benchmark with scBasset

scBasset ^9^ is a sequence-based CNN method to model single-cell ATAC-seq data. We followed the guidelines on the scBasset GitHub repository at https://github.com/calico/scBasset. scBasset takes a scATAC-seq anndata file and a genome fasta file as input. The genome fasta file can be downloaded from https://hgdownload.soe.ucsc.edu/downloads.html. We used the command-line tool ‘scBasset/bin/scbasset_preprocess.py’ to extract sequences underlying peak regions, one-hot encode them, and save them in sparse h5 format. The model was trained using ’scBasset/bin/scbasset_train.py’, and model selection was based on training auROC. Default training parameters were adopted: --bottleneck 32, --batch_size 128, --lr 0.01, --epochs 1000. In the within-sample scenario, we used the predict() function to obtain the model’s predictions on the test set and calculate auROC and auPRC. We applied the imputation_Y_normalize() function to obtain the model’s denoised (sequencing depth-normalized) results, and computed the NS and LS metrics on the denoised matrix.

### Benchmark with Seurat

To predict chromatin accessibility, we followed the guidelines on the Seurat (v.4.4.0) ^5^ website at https://satijalab.org/seurat/archive/v3.2/integration.html. We used the FindTransferAnchors() function with the parameter: reduction = ‘cca’ to build anchors between training and testing scRNA-seq data, choosing the first 30 CCA dimensions (dims = 1:30). We then applied the TransferData() function to transfer scATAC-seq data from training set to test set using the found anchors, with k.weight = 10 specifying the number of KNNs. In the within-sample scenario, when visualizing the prediction results using UMAP, Seurat can only make predictions for the test data. Therefore, we used raw scATAC-seq data to supplement and construct a complete dataset as the plot background, but the raw scATAC-seq data were not involved in metric calculations. For each sample, we maintained the same training and test set divisions as in XChrom to ensure consistency in evaluation.

### Benchmark with LS_Lab

We followed the tutorial provided on the neurips2021_multimodal_topmethods GitHub repository: https://github.com/openproblems-bio/neurips2021_multimodal_topmethods/blob/main/src/predict_modality/methods/LS_lab/run/script.py/. For the “gex2atac” task, LS_Lab ^6^ uses a KNN regression model to predict scATAC-seq data from scRNA-seq data through 10-fold cross-validation. We first performed dimensionality reduction on both scRNA-seq and scATAC-seq data using the TruncatedSVD() function with n_components = 50, and then applied KNeighborsRegressor() with parameters n_neighbors = 25 and metric = ’minkowski’. This model learned the mapping between the two modalities in the latent space. The predictions in the latent space were projected back into the original feature space by multiplying them with the component matrix from the dimensionality reduction model. The final prediction was obtained by averaging the 10-fold cross-validation results ^4^. For each sample, we maintained the same training and test set divisions as in XChrom to ensure consistency in evaluation.

### Benchmark with MultiVI

We followed the guidelines provided on the MultiVI ^7^ website at https://docs.scvi-tools.org/en/stable/tutorials/notebooks/multimodal/MultiVI_tutorial.html. We used the get_accessibility_estimates() function of MultiVI to predict chromatin accessibility from scRNA-seq data. All parameters were kept at default settings. For each sample, we maintained the same training and test set divisions as in XChrom to ensure consistency in evaluation.

### Identification of cell-type-specific motifs using ISM

For the human-specific chromatin accessibility regions (lacking homology to the mouse genome) near the *PSEN1* and *DNAJC5* genes, we performed ISM to compute single-nucleotide importance scores at single-cell resolution. The process was as follows: For each target sequence position, we performed three forward passes using the mouse-trained XChrom model, mutating each reference nucleotide to one of the three alternative nucleotides while keeping the initial human-derived cell embeddings unchanged as the model input. By comparing the chromatin accessibility predictions for each mutated sequence with the reference sequence, we computed the effect of each mutation in each cell. The ISM scores for the four nucleotides at each position were normalized such that their sum was zero, and the normalized ISM score for the reference nucleotide was used as the importance score for that position. This process yielded a complete ISM score matrix for each target peak region, with dimensions (cell number × 1344 bp × 4 nucleotides), quantifying the impact of nucleotide mutations at each position on the predicted values for each cell. For further analysis, we performed *Z*-score normalization of the ISM scores across different cells to enable comparison of the relative importance of each nucleotide position across cell types. Additionally, we used the FIMO tool ^26^ to scan the target peak regions for TF motifs, setting a candidate motif matching *P*-value threshold to 1 × 10□^3^. For each identified candidate motif, we computed the dot product between its position weight matrix (PWM) and the corresponding ISM score for the motif region. This yielded a specificity score for each cell. We also performed a two-sided Wilcoxon rank-sum test to evaluate statistical differences between the specificity scores of any two cell types.

### Calculate TF motif activity

To quantitatively assess TF regulatory activity at the single-cell level, we adopted a motif insertion-based perturbation strategy and used the XChrom model to calculate motif activity score for each TF in every cell. First, we applied the fasta_dinucleotide_shuffle tool ^58^ to randomly select and shuffle the sequences of 1000 peak regions from the training/test peak sets, generating 1000 background genome sequences as controls. In the cross-species analysis, the background sequences were derived from peak regions in the test set; for the COVID-19 analysis, the background sequences were constructed using the peak regions from the training set (PBMC10k).

For each TF in the motif database, we sampled 1000 representative motif sequences from its PWM, each of which was inserted into the central position of a corresponding background sequence, resulting in 1000 motif-inserted sequences for each TF motif. We then input the motif-inserted sequences and the original background sequences into the trained XChrom model for forward propagation, using initial cell embeddings derived from the scRNA-seq data of the test set as input, and predicted sequencing depth-normalized chromatin accessibility values for all cells. The difference in prediction between the motif-inserted sequences and the background sequences was defined as the motif activity score in each cell. For each TF, we calculated the mean motif activity score across all 1000 sequences in each cell, obtaining the raw TF activity score for that cell. To visualize the activity distribution of a specific TF across different cells, we performed *Z*-score normalization on the raw TF activity scores for all cells to facilitate comparison.

We used CIS-BP ^59^ as the motif database in this analysis, which was downloaded from https://meme-suite.org/meme/meme-software/Databases/motifs/motif_databases.12.27.tgz. For human samples as the test set, we used the human motif database (motif_databases/CIS-BP_2.00/Homo_sapiens.meme), and for mouse samples as the test set, we used the mouse motif database (motif_databases/CIS-BP_2.00/Mus_musculus.meme).

### TF activity correlation between species across cell types

To assess the conservation of TF regulatory activity between species, we calculated the Pearson correlations of TF motif activity across cell types under three different settings:

(i) Correlation analysis using a human-trained model: We used the XChrom model trained on human data to calculate TF motif activity in both human (m1d1) and mouse (mop3c2) samples, and then calculated the Pearson correlation coefficient of each TF’s activity between human and mouse samples across cell types.
(ii) Correlation analysis using a mouse-trained model: We used the XChrom model trained on mouse data to calculate TF motif activity in both human (m1d1) and mouse (mop3c2) samples, and then calculated the Pearson correlation coefficient of each TF’s activity between human and mouse samples across cell types.
(iii) Correlation analysis between models from different species: We calculated TF motif activity using the human-trained model in human sample (m1d1) and the mouse-trained model in mouse sample (mop3c2), and analyzed the correlation across cell types between the two species.

For each cell type, we computed the mean activity value across all cells. All analyses were based on 677 common motifs shared between the human and mouse motif databases to ensure consistency in the comparison.

### Annotation of PBMC10k training data

We used the reference mapping method from the Seurat (v4.4.0) package ^5^ (https://satijalab.org/seurat/articles/multimodal_reference_mapping#intro-seurat-v4-reference-mapping) to annotate cell types in the PBMC10k scRNA-seq data. First, we loaded an annotated multimodal PBMC dataset (pbmc_multimodal.h5seurat from https://atlas.fredhutch.org/nygc/multimodal-pbmc/) as a reference. After performing SCTransform() normalization on the query data (PBMC10k), we used the FindTransferAnchors() function (parameters: normalization.method = “SCT”, reference.reduction = “spca”, dims = 1:50) to establish cell correspondences between the reference and query data. Then, we applied the MapQuery() function (parameters: reference.reduction = “spca”, reduction.model = “wnn.umap”) to transfer cell type labels from the reference (including celltype.l1 and celltype.l2 labels) to the query data, followed by UMAP visualization using the reference’s “ref.umap” coordinates. Finally, the annotated query data was saved in both h5Seurat and h5ad formats for further analysis.

Following initial annotation, we further refined monocyte subcluster annotations within PBMC10k. Using the processed COVID-19 scRNA-seq dataset (COVID19_MHH50/scRNA/Monocytes_scRNAseq.rds, including PCA reduction and UMAP visualization results) as a reference, we implemented a PCA-based reference mapping method for cell type annotation. Specifically, after extracting monocytes from the PBMC10k dataset, we normalized the data using LogNormalize(), identified the top 1000 highly variable genes, and scaled the data. We then used the FindTransferAnchors() function (parameters: reference.reduction = “pca”, dims = 1:20) to establish transcriptomic similarity between the reference and query monocyte data. Finally, we mapped the reference’s cell type labels (new.id: R1-R8) to the query dataset with MapQuery() function (parameters: reference.reduction = “pca”, reduction.model = “umap”) and performed projection using the reference’s UMAP model.

### Differential analysis of TF activity in the COVID-19 dataset

Using an XChrom model trained on PBMC10k, we predicted the TF activities for cells in the COVID-19 scRNA-seq dataset. Specifically, we extracted monocytes from the COVID-19 TF activity matrix and calculated the mean differences in TF activity between R1 (4,271 cells) and other subclusters (34,178 cells). We used the two-sided Wilcoxon rank-sum test, with *P*-value adjustment via the false discovery rate (FDR) correction procedure (Benjamini-Hochberg method). The same analysis was performed for R4 (7,100 cells) versus other subclusters (31,349 cells).

Furthermore, we analyzed the mean differences in TF activity among cMono between the severe group (18,463 cells) and the mild group (18,947 cells). Inter-group comparisons were performed using the two-sided Wilcoxon rank-sum test, with *P*-value adjustment via FDR correction. We visualized the results using a volcano plot (Fig. 6d), with mean differences on the x-axis and adjusted *P*-values on the y-axis. The x-axis was scaled using a pseudo-log transformation from the scales package (scale_x_continuous(trans = scales::pseudo_log_trans(base = 3, sigma = 0.001), breaks = scales::pretty_breaks(n = 3))). Additionally, we extracted the expression matrix of cMono from the raw scRNA-seq data and normalized it using scanpy.pp.normalize_total() and scanpy.pp.log1p() ^56^. We calculated the Pearson correlation coefficients between activity and expression values across cells for 863 common TFs, with *P*-values adjusted for multiple comparisons using FDR correction.

### Motif enrichment analysis in COVID-19 scATAC-seq data

We used processed COVID-19 scATAC-seq dataset for further analysis. For monocyte subclusters (C1-C6), we used the getMarkerFeatures() function from the ArchR ^60^ tool (parameters: useMatrix = “PeakMatrix”, groupBy = “Clusters”, bias = c(“TSSEnrichment”, “log10(nFrags)”), testMethod = “wilcoxon”) to identify marker peaks for each subcluster. We then applied the peakAnnoEnrichment() function (parameters: peakAnnotation = “Motif”, cutOff = “Pval <= 0.05 & Log2FC > 1”) to identify significantly enriched TF motifs in these marker peaks. Motif annotations were based on the CIS-BP database ^59^. According to the original study, the C1 and C4 subclusters in the scATAC-seq data were mapped to the R1 and R4 subclusters in the scRNA-seq data, respectively ^34^.

## Data availability

In the within-sample scenario, we downloaded the 10x Multiome datasets from 10x Genomics: human PBMC dataset (h_PBMC) https://support.10xgenomics.com/single-cell-multiome-atac-gex/datasets/2.0.0/pbmc_granulocyte_sorted_3k, mouse brain dataset (m_brain) https://support.10xgenomics.com/single-cell-multiome-atac-gex/datasets/2.0.0/e18_mouse_brain_fresh_5k, and human healthy brain dataset (h_brain) https://www.10xgenomics.com/datasets/frozen-human-healthy-brain-tissue-3-k-1-standard-2-0-0.

The 8.6-week female gonad dataset (h_gonads) is available from ArrayExpress (https://www.ebi.ac.uk/biostudies/arrayexpress), with accession number E-MTAB-11708 and sample name HCA_F_GON10535495 ^61^. The mouse secondary palate at embryonic day 12.5 dataset (m_palates) is available from Gene Expression Omnibus (GEO, https://www.ncbi.nlm.nih.gov/geo/) under accession code <u>GSE218576</u> ^62^. In the cross-sample scenario, we used human BMMC datasets from GEO under accession code <u>GSE194122</u> ^17^. In the cross-species scenario, the 10x Multiome datasets for the motor cortex of human, mouse, macaque, and marmoset are available from GEO under accession code <u>GSE229169</u> ^21^. We downloaded the COVID-19 datasets ^34^ from https://nubes.helmholtz-berlin.de/s/wqg6tmX4fW7pci5/download. In this study, processed data after batch effect correction and ready for model training have been uploaded to Zenodo: https://doi.org/10.5281/zenodo.16959682.

## Code availability

XChrom is freely available as an open-source Python package at https://github.com/Miaoyuanyuan777/XChrom. Original codes and scripts used for the analysis are available at https://github.com/Miaoyuanyuan777/XChrom_analysis. Tutorials are provided at https://xchrom.readthedocs.io/en/latest/index.html, covering XChrom’s installation, training, evaluation, and interpretation under various scenarios described in this study.

## Acknowledgements

This work was supported by the National Natural Science Foundation of China (32370690), the Major Project of Guangzhou National Laboratory (GZNL2024A01003), and the National Key R&D Program of China (2021YFA1100501). We thank Prof. Sijia Wang for helpful discussions on this work.

## Author Contributions

Y.M. and Z.W. designed the study. Y.M. performed data analysis. Y.M. built the software. X.L. and W.Z. tested the software. Y.M. wrote the manuscript. Z.W., X.L., D.H., Y.L., and W.Z. revised the manuscript. Z.W. supervised the study.

