## Supplementary Figures for "XChrom: a cross-cell chromatin accessibility prediction model integrating genomic sequence and cellular context"

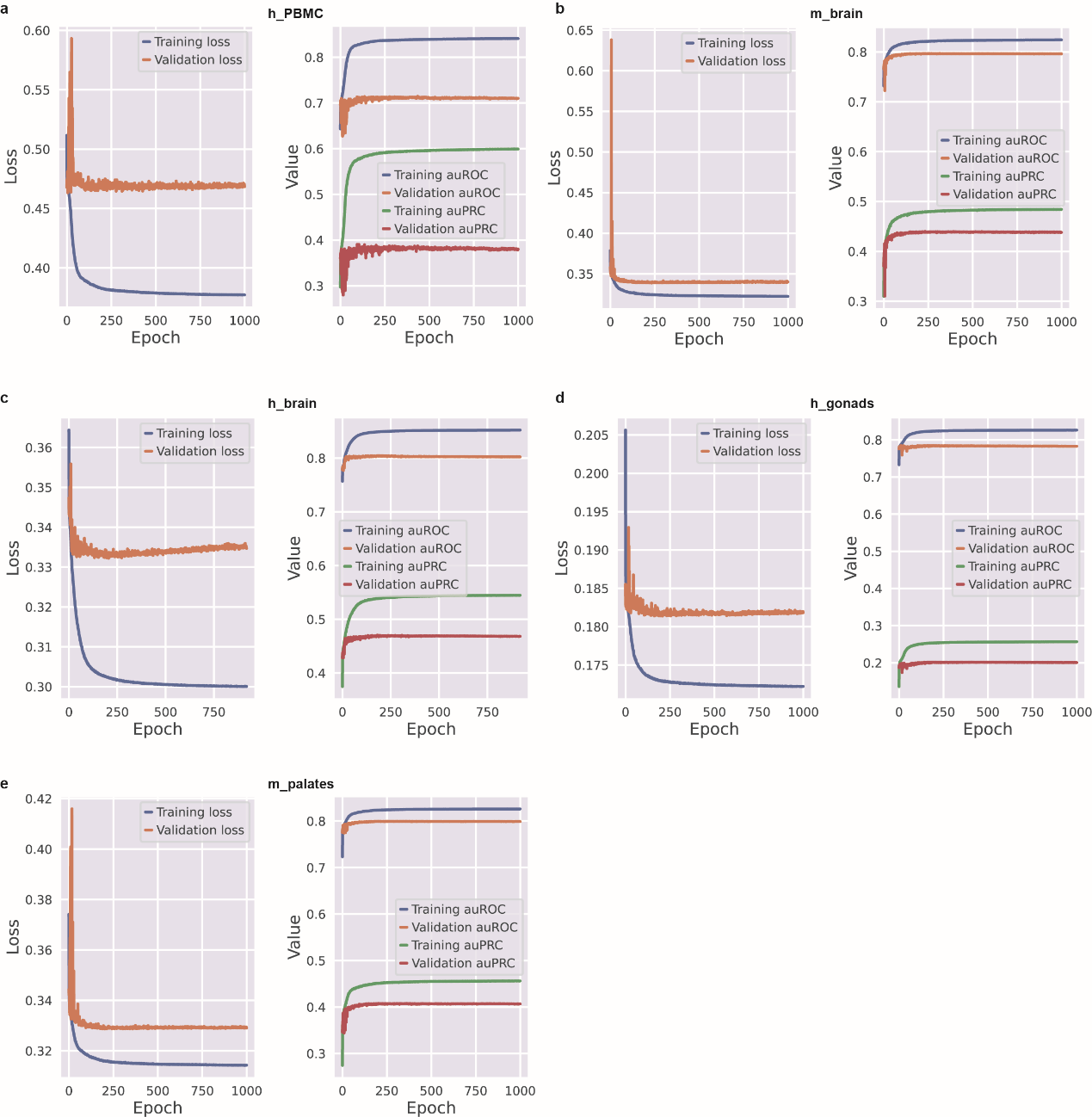

**Supplementary Fig. 1: Training monitoring curves across datasets in the within-sample scenario.** For each dataset, the left panel shows training and validation loss per epoch; the right panel shows training and validation auROC and auPRC per epoch. (a) h_PBMC; (b) m_brain; (c) h_brain; (d) h_gonads; (e) m_palates.

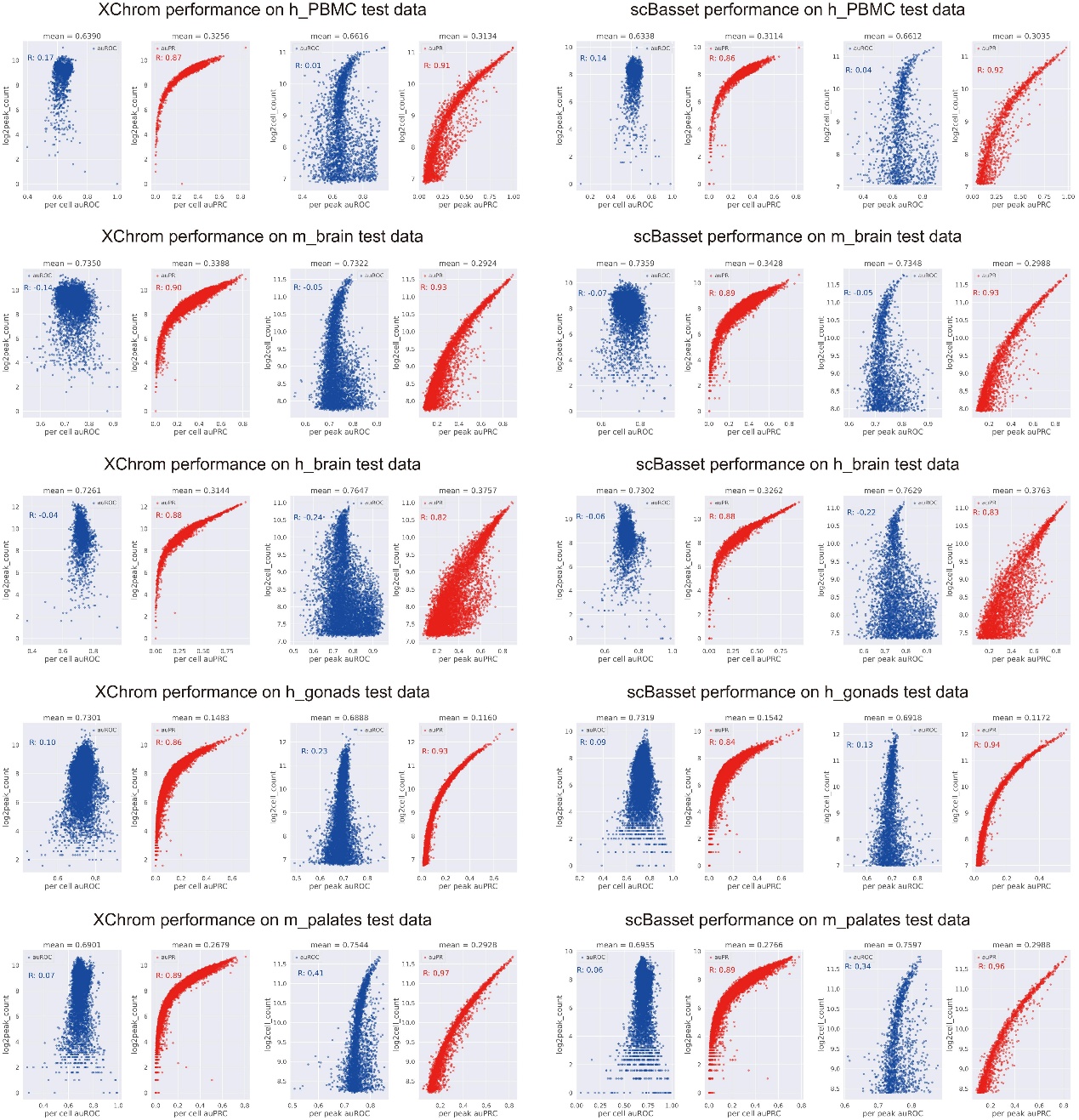

**Supplementary Fig. 2: Cross-region prediction performance of XChrom and scBasset on a single sample.** Rows correspond to datasets (top to bottom is h_PBMC, m_brain, h_brain, h_gonads, m_palates); columns to models (left panel: XChrom, right panel: scBasset). Within each panel, the four subplots (left to right) show: (i) auROC per cell vs. log-transformed peak counts per cell; (ii) auPRC per cell vs. log-transformed peak counts per cell; (iii) auROC per peak vs. log-transformed cell counts per peak; and (iv) auPRC per peak vs. log-transformed cell counts per peak.

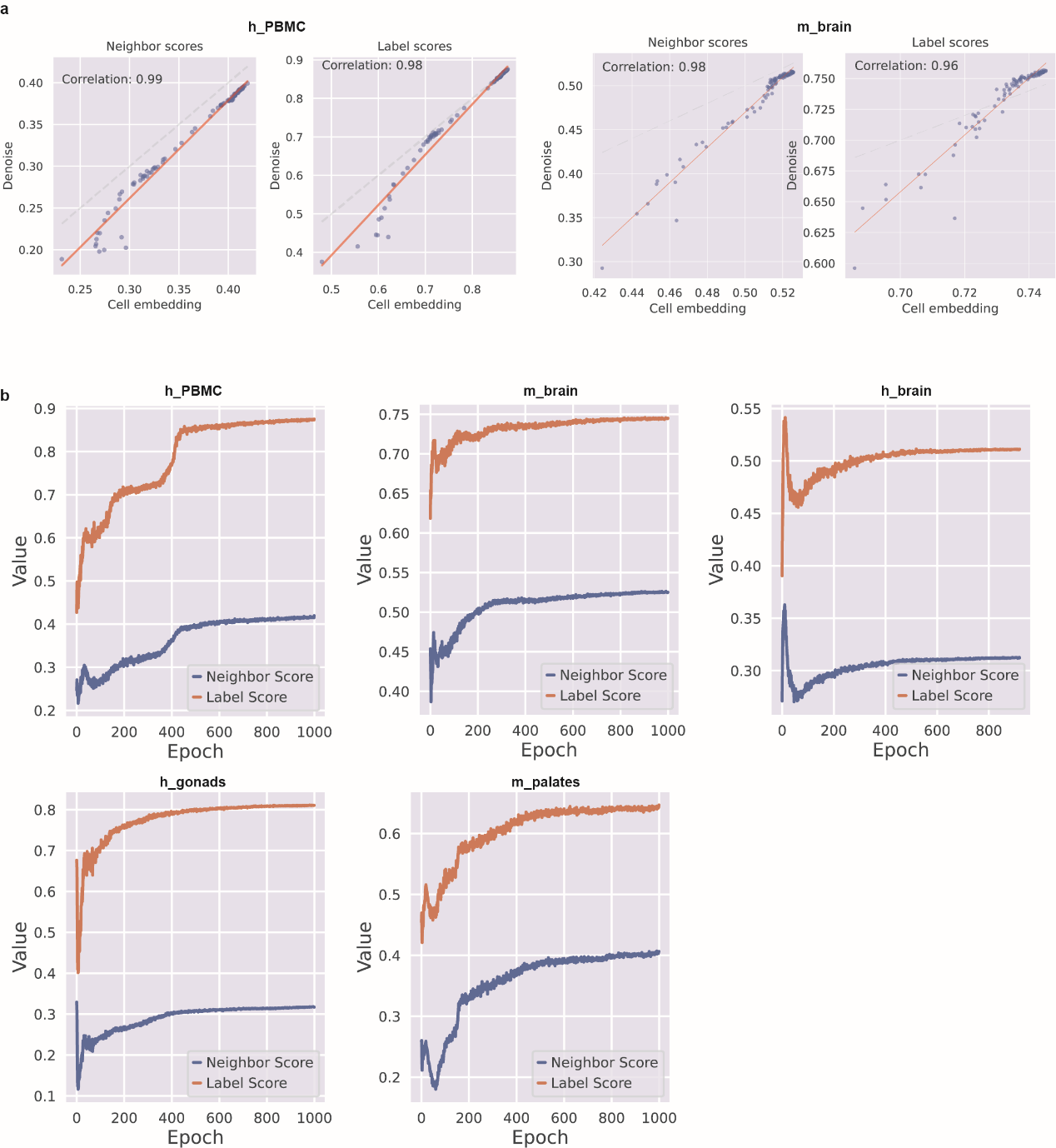

**Supplementary Fig. 3: Monitoring the XChrom training process using NS (*k* = 100) and LS (*k* = 100).**

a, For XChrom trained on h_PBMC and m_brain, Pearson correlation between metrics computed from the final cell embeddings and from the model-denoised scATAC-seq profiles. The left pair of subplots corresponds to h_PBMC and the right pair to m_brain; within each pair, the left subplot shows Pearson correlation of NS (*k* = 100) and the right subplot shows LS (*k* = 100).

b, NS (*k* = 100) and LS (*k* = 100) computed from the cell embeddings at each epoch during training, shown across five datasets.

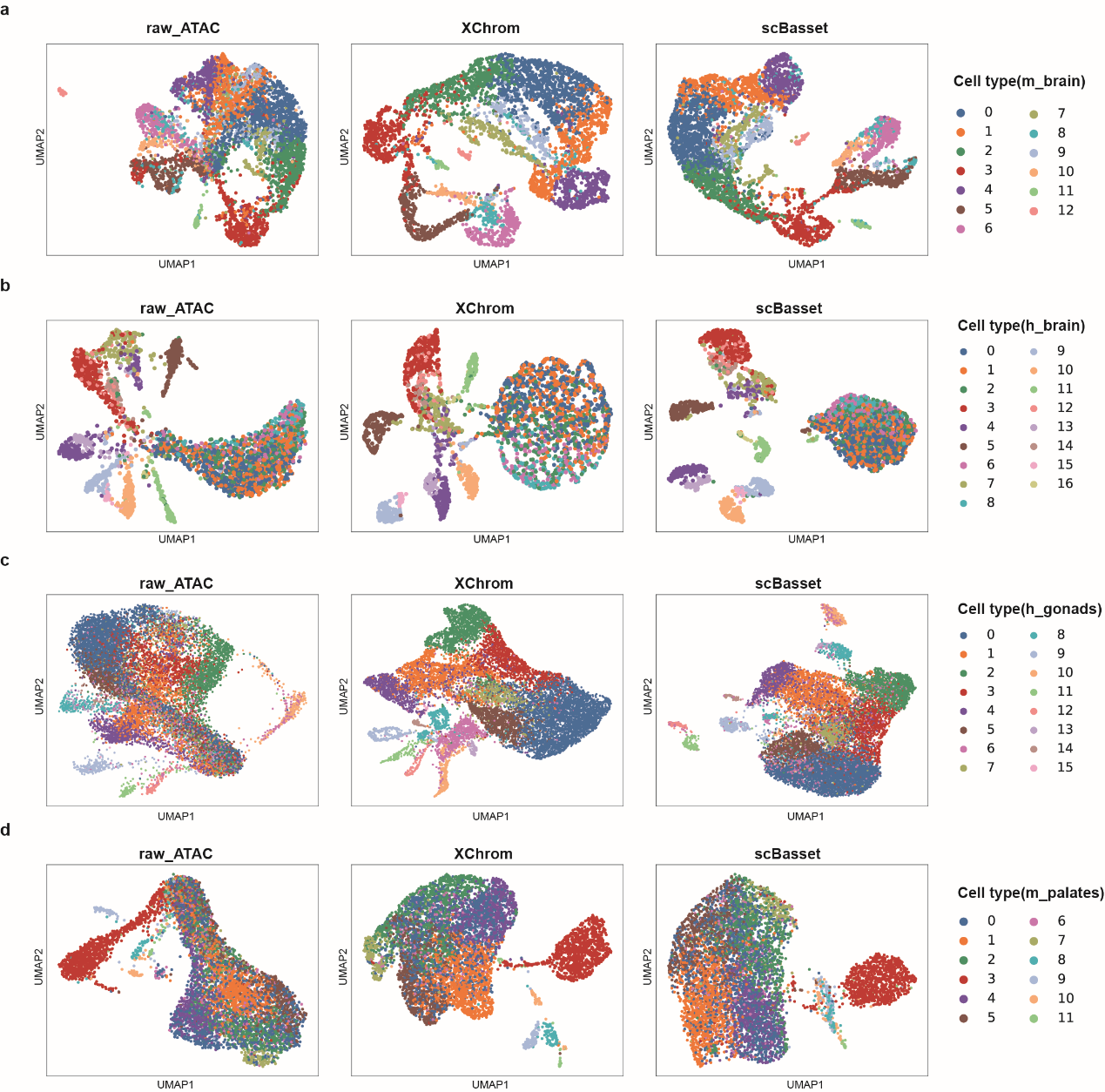

**Supplementary Fig. 4:** UMAP visualizations of raw scATAC-seq data and of the whole dataset denoised by XChrom and scBasset. Panels: (a) m_brain; (b) h_brain; (c) h_gonads; (d) m_palates.

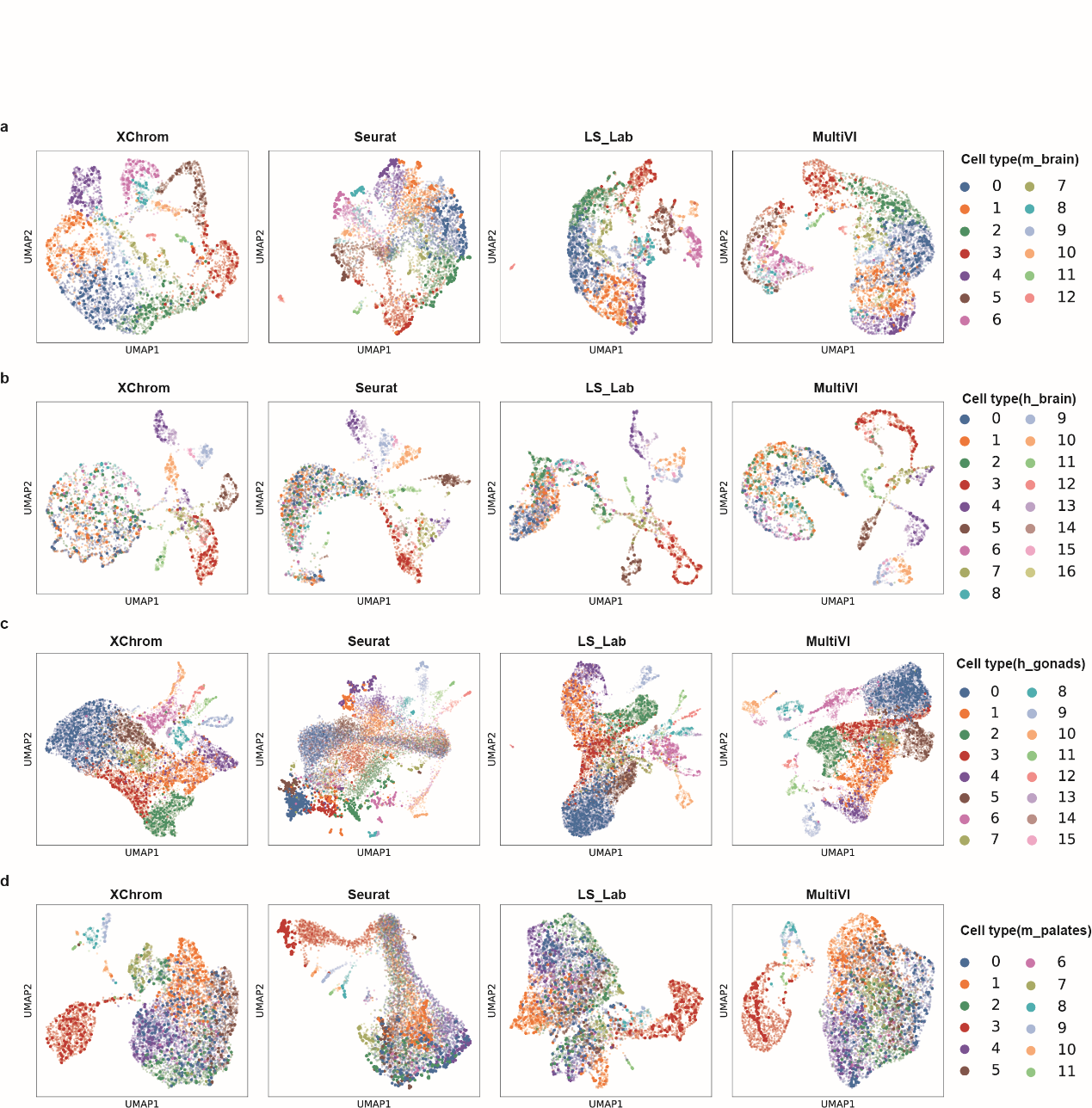

**Supplementary Fig. 5:** UMAP visualizations of test cell predictions by XChrom, Seurat, LS_Lab, and MultiVI across datasets. m_brain (a); h_brain (b); h_gonads (c); m_palates (d). Larger points denote test cells, while background points denote training cells.

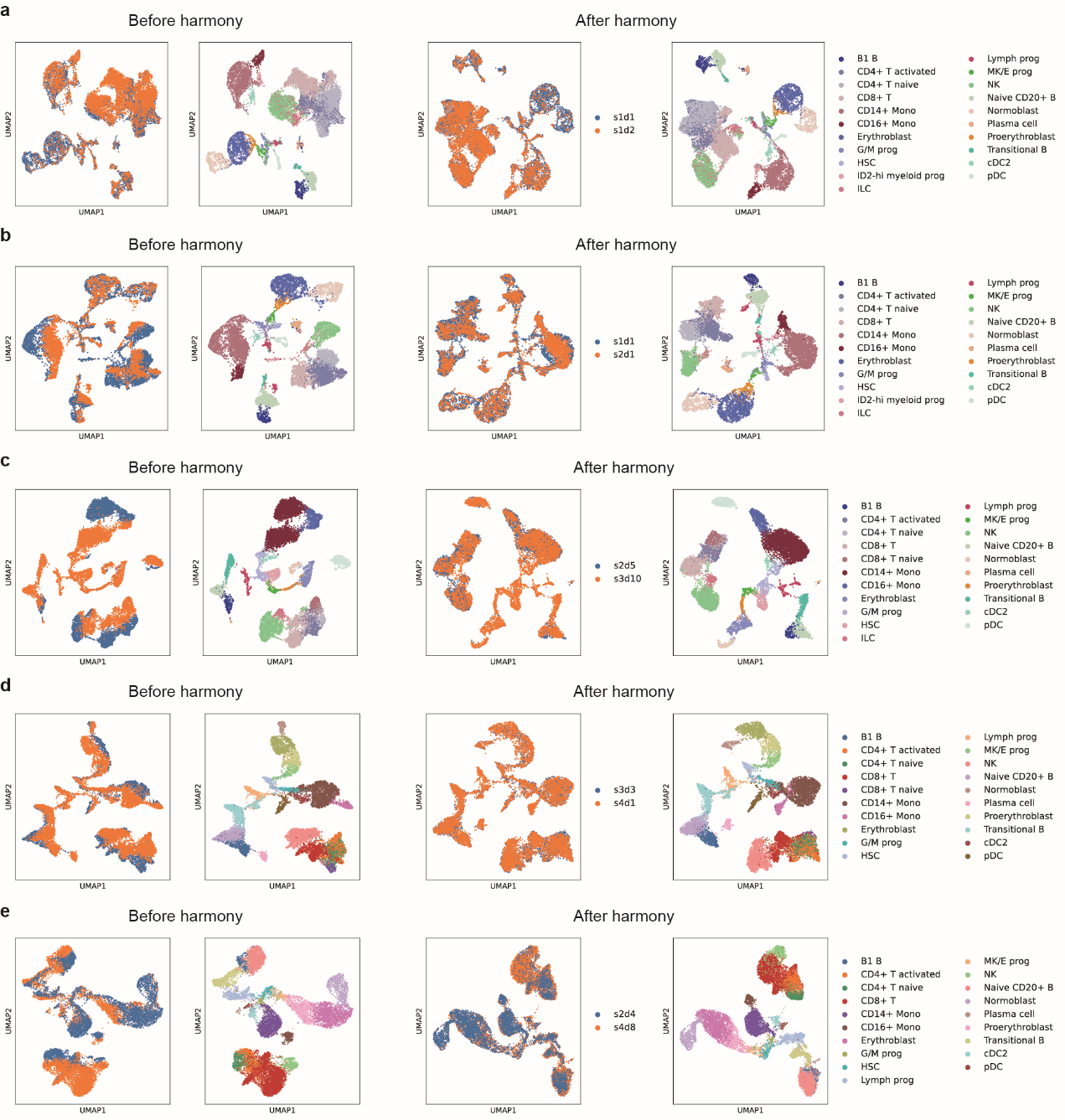

**Supplementary Fig. 6:** **Batch-effect correction in the cross-sample scenario with Harmony.** UMAP visualizations of the training and test sets before and after Harmony-based batch correction (treating each sample as a batch). For each panel, columns show before Harmony correction (left) and after Harmony correction (right). Panel titles label the ‘training set—test set’ sample pair. Panels: (a) s1d1-s1d2; (b) s1d1-s2d1; (c) s2d5-s3d10; (d) s3d3-s4d1; (e) s2d4-s4d8.

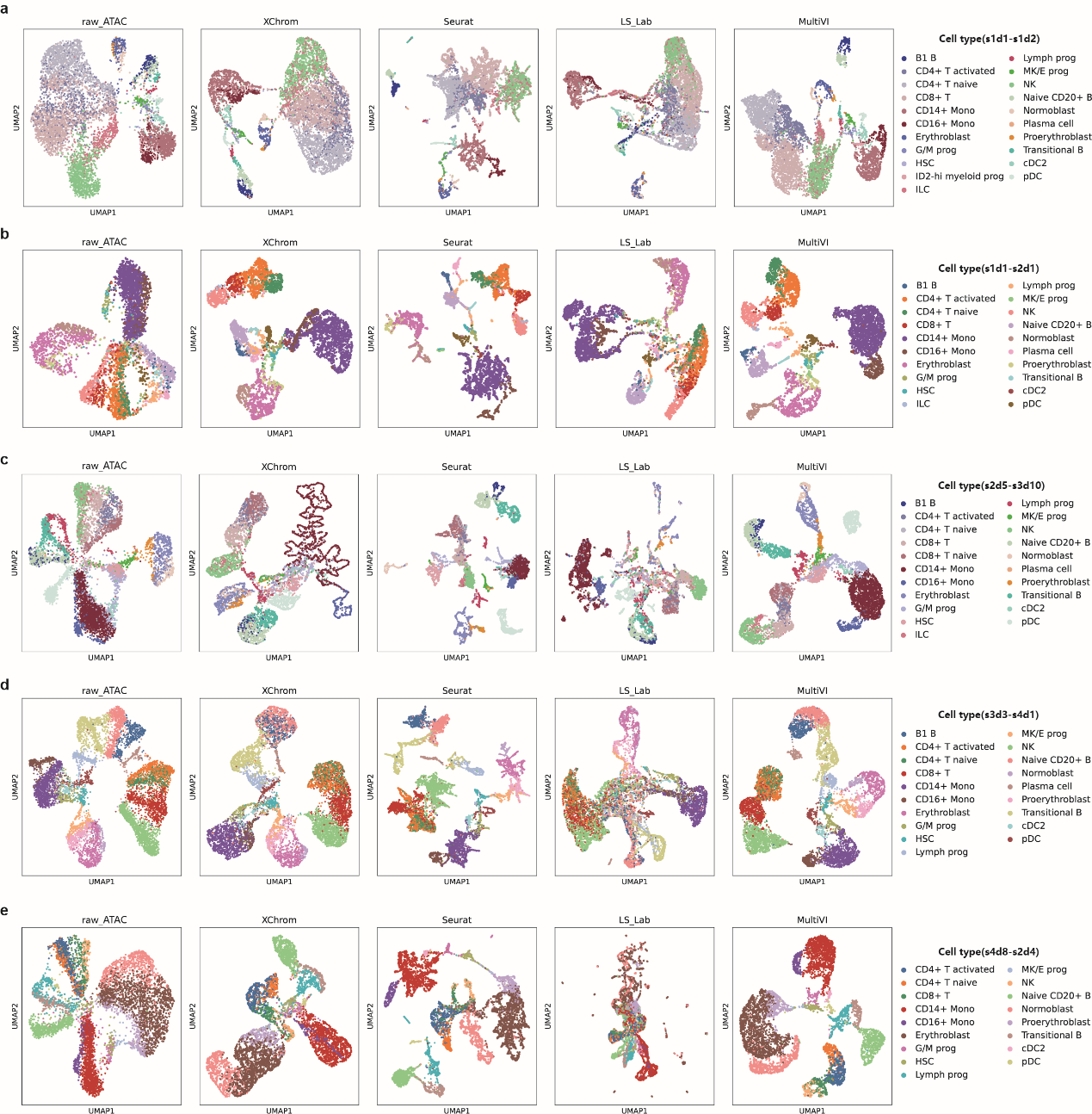

**Supplementary Fig. 7: Cross-sample (one-to-one) prediction.** UMAP visualizations of raw scATAC-seq and predictions from XChrom, Seurat, LS_Lab, and MultiVI for each training–test sample pair. Columns (left to right): raw_ATAC, XChrom, Seurat, LS_Lab, MultiVI. Panel titles indicate the training–test sample pair. Panels: (a) s1d1-s1d2; (b) s1d1-s2d1; (c) s2d5-s3d10; (d) s3d3-s4d1; (e) s2d4-s4d8.

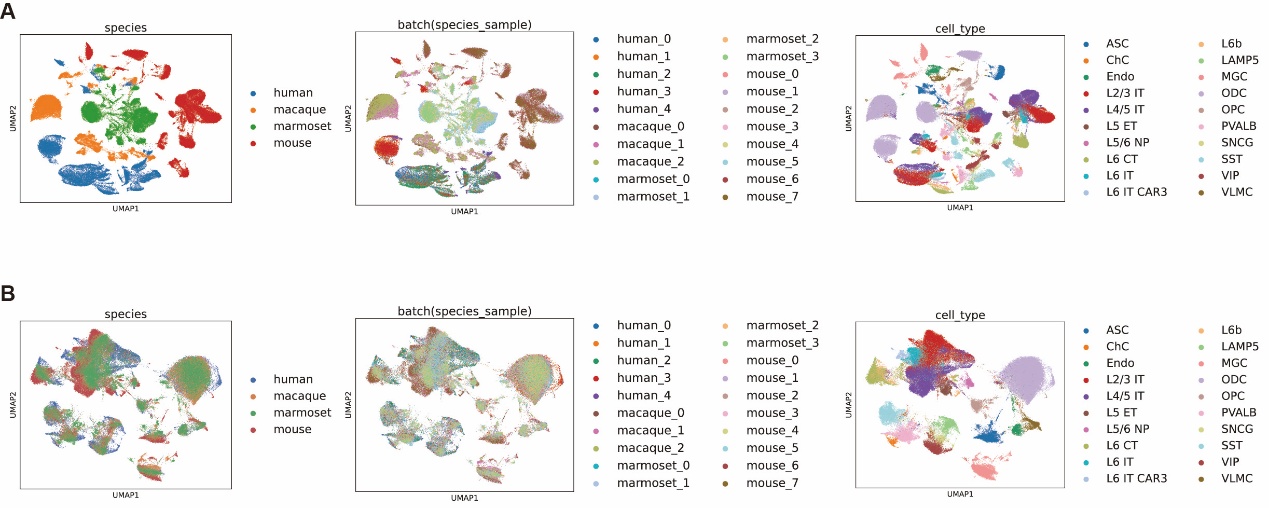

**Supplementary Fig. 8:** **Batch-effect correction in the cross-species scenario with harmony.** UMAP visualizations of the different species samples before and after Harmony-based batch correction (treating each sample as a batch). Before batch correction (a) and after batch correction (b). For each panel, left to right, colored with species, samples, and cell types.

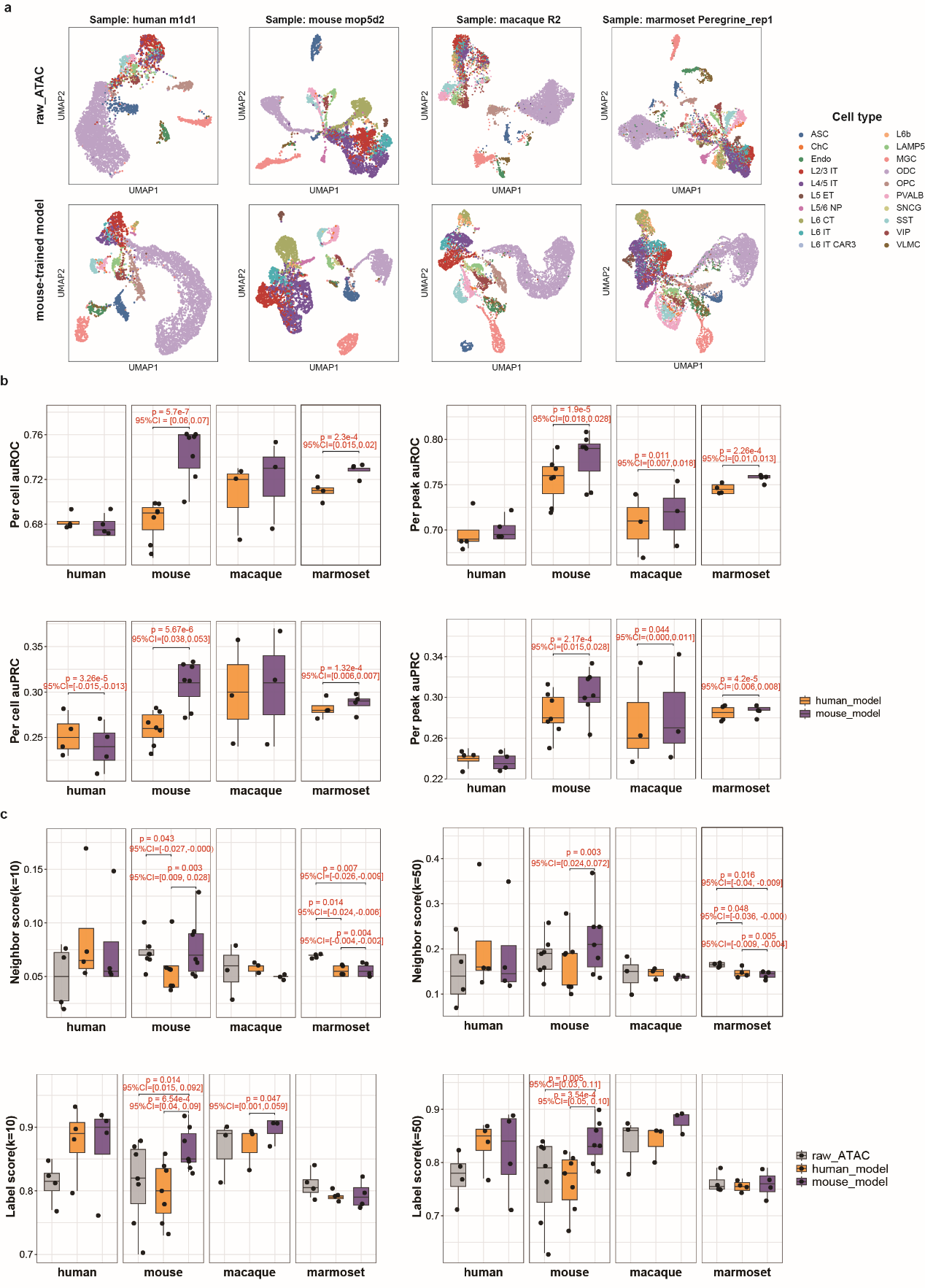

**Supplementary Fig. 9:** **XChrom’s prediction performance in cross-species scenario.**

a, UMAP visualizations of predictions generated by the mouse_model when applied to the test sets from four different species. Sample names are labelled in the plot.

b, Comparison of model performance between human_model and mouse_model using per cell auROC/auPRC (left) and per peak auROC/auPRC (right) values computed from predictions on all samples. *P*-values and 95% CIs are shown in the figure if *P*-value<0.05 (two-sided paired t-test).

c, Comparison of model performance between human_model and mouse_model using NS/LS (*k*=10) (left) and NS/LS (*k*=50) (right) values computed from predictions on all samples. *P*-values and 95% CIs are shown in the figure if *P*-value<0.05 (two-sided paired t-test).

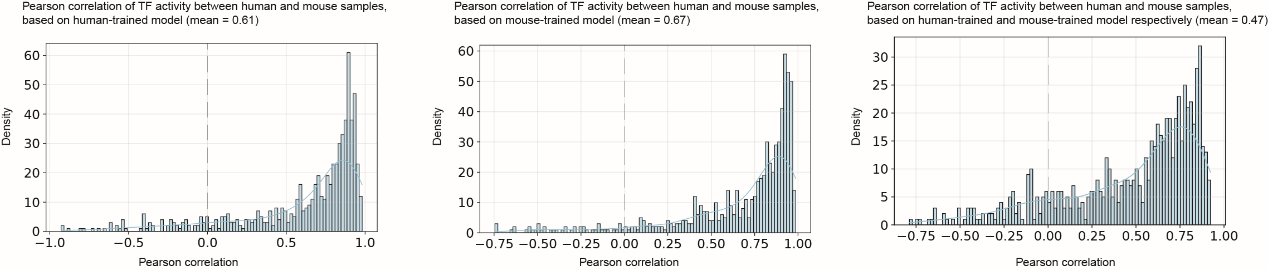

**Supplementary Fig. 10:** **Cross-species correlations of TF-motif activity.** Pearson correlations between human and mouse across cell types, using the 677 motifs shared between the human and mouse motif databases. Panels (left to right): left, human_model predictions for both human and mouse samples; middle, mouse_model predictions for both human and mouse samples; right, human_model predictions for the human sample and mouse_model predictions for the mouse sample.

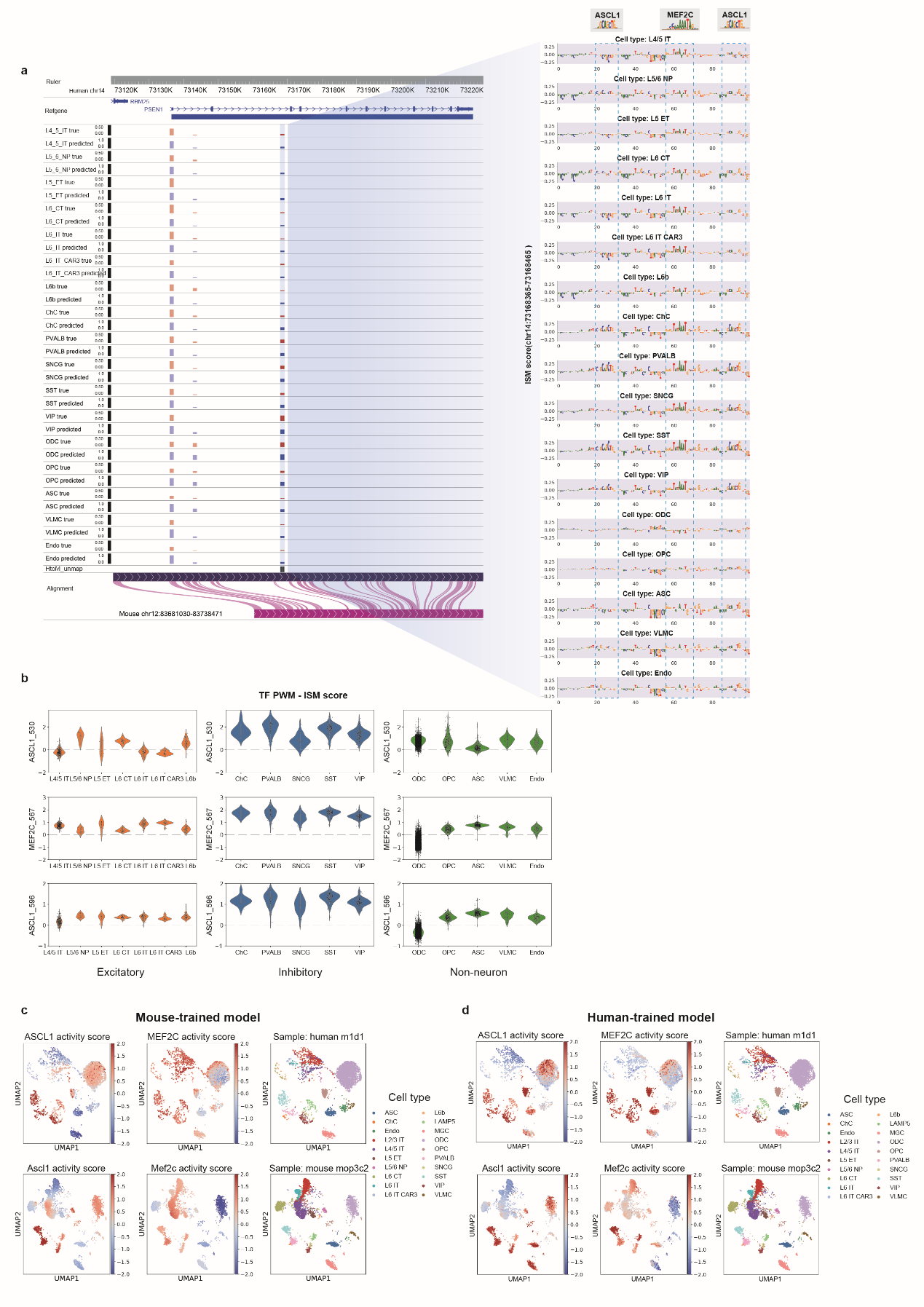

**Supplementary Fig. 11:** **XChrom interpretability analysis using the *PSEN1* regulatory sequence in cross-species scenario.**

a, Visualization of mouse_model predictions for a *PSEN1* regulatory sequence in the human sample (m1d1), with reference data from the WashU Comparative Epigenome Browser (https://epigenome.wustl.edu/BrainComparativeEpigenome/). The blue-shaded sequence represents the human region (chr14: 73167855-73169199) that lacks homology to the mouse genome. Subsequences matching MEF2C and ASCL1 motifs are highlighted with blue dashed boxes in the cell-type-averaged ISM score plot.

b, Per-cell PWM-ISM scores for MEF2C and ASCL1 in excitatory (including L4/5 IT, L5/6 NP, L5 ET, L6 IT, L6 IT CAR3, L6b cell types), inhibitory (including ChC, PVALB, SNCG, SST, VIP cell types), and non-neuron (including ODC, OPC, ASC, VLMC, Endo cell types). The PWM-ISM score is the dot product of the PWM and ISM scores at motif match sites (MEF2C at chr14:73168422-73168436, ASCL1 at chr14:73168385-73168397 and chr14:73168451-73168463).

c-d, UMAP visualizations of MEF2C and ASCL1 motif activity predicted by XChrom across different cell types in the human sample (top) and the mouse sample (bottom). XChrom was trained on the mouse sample (mouse_model) (c) and the human sample (human_model) (d).

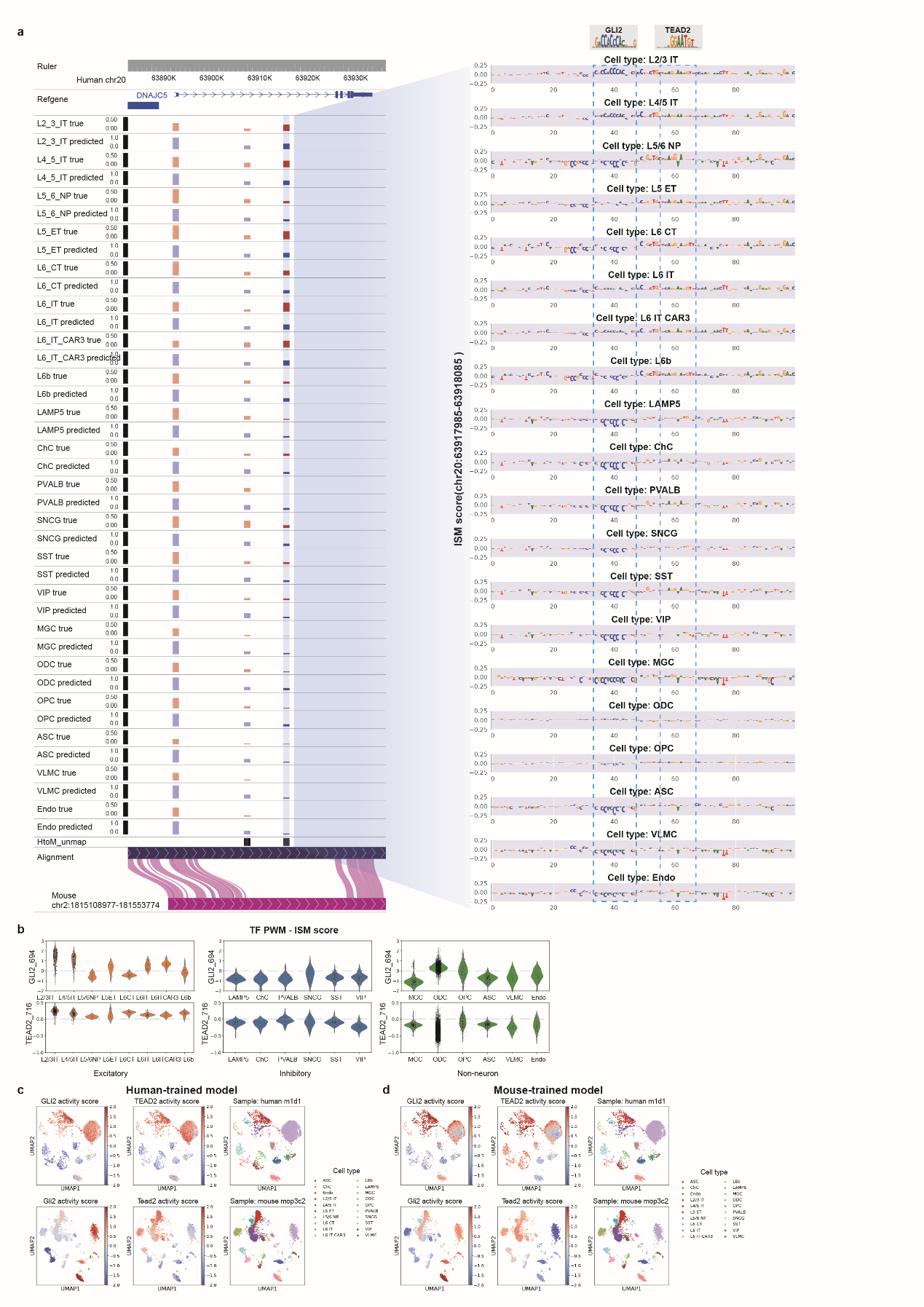

**Supplementary Fig. 12:** **XChrom interpretability analysis using the *DNAJC5* regulatory sequence in cross-species scenario.**

a, Visualization of mouse_model predictions for a *DNAJC5* regulatory sequence in the human sample (m1d1), with reference data from the WashU Comparative Epigenome Browser (https://epigenome.wustl.edu/BrainComparativeEpigenome/). The blue-shaded sequence represents the human region (chr20: 63917325-63918669) that lacks homology to the mouse genome. Subsequences matching GLI2 and TEAD2 motifs are highlighted with blue dashed boxes in the cell-type-averaged ISM score plot.

b, Per-cell PWM-ISM scores for GLI2 and TEAD2 in excitatory (including L2/3 IT, L4/5 IT, L5/6 NP, L5 ET, L6 IT, L6 IT CAR3, L6b cell types), inhibitory (including LAMP5, ChC, PVALB, SNCG, SST, VIP cell types), and non-neuron (including MGC, ODC, OPC, ASC, VLMC, Endo cell types). The PWM-ISM score is the dot product of the PWM and ISM scores at motif match sites (GLI2 at chr20:63918019-63918033, TEAD2 at chr20: 63918041-63918053).

c-d, UMAP visualizations of GLI2 and TEAD2 motif activity predicted by XChrom across different cell types in the human sample (top) and the mouse sample (bottom). XChrom was trained on the human sample (human_model) (c) and the mouse sample (mouse_model) (d).

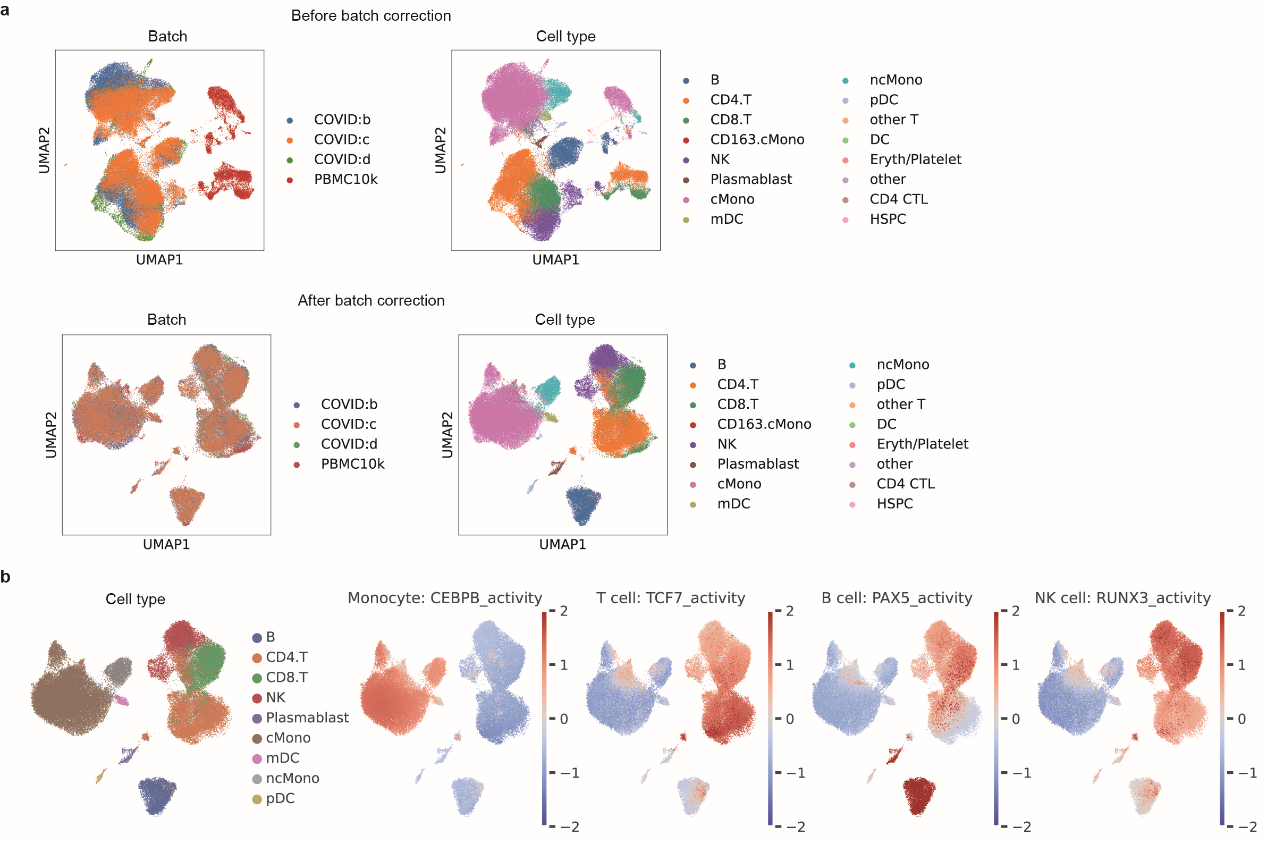

**Supplementary Fig. 13:** **Application of XChrom to PBMC datasets from COVID-19 patients.**

a, UMAP visualizations of the PBMC10k and COVID-19 scRNA-seq datasets before and after Harmony-based batch correction. Rows: before correction (top), after correction (bottom). Within each panel: left subplot colored by batch; right subplot colored by cell type.

b, UMAP visualizations colored by predicted motif activity for CEBPB, TCF7, PAX5, and RUNX3 from a PBMC10k-trained XChrom model applied to the COVID-19 dataset. For visualization, activity scores for each TF motif were *Z*-score normalized across all cells. Panels (left to right): CEBPB, TCF7, PAX5, RUNX3.
